## Supplemental text for "Inferring selection effects in SARS-CoV-2 with Bayesian Viral Allele Selection"

This supplement is organized as follows. In Sec. S1 we describe the discrete-time branching process and diffusion approximation that forms the basis of our modeling approach. In Sec. S2 we summarize the simplest approach to statistical inference that utilizes the resulting diffusion-based likelihood, namely maximum a posteriori (MAP) inference. In Sec. S3 we describe our approach to inducing sparsity in the same class of diffusion-based models via Bayesian Variable Selection. In Sec. S4 we give a high-level overview of how we do Markov Chain Monte Carlo (MCMC) inference in this class of models. In Sec. S5 we generalize the discussion to the case of multiple spatial regions. In Sec. S6 we introduce the effective population size estimators we use. In Sec. S7 we generalize our basic model to include vaccination-dependent selection effects. In Sec. S8 we discuss the effect of sampling, i.e. the fact that not all viral sequences are observed. In Sec. S9 we provide an extended discussion of the prior inclusion probability $h$, which is one of the main hyperparameters of our method. In Sec. S10 we discuss alternative approaches to modeling and inference and why we prefer Bayesian Viral Allele Selection. In Sec. S11 we briefly discuss how we report uncertainty estimates. In Sec. S12 we discuss some of the limitations of BVAS. In Sec. S13 we describe how we simulate viral dynamics and present additional experiments not included in the main text. In Sec. S14 we give additional details and report additional results for our SARS-CoV-2 analysis. In Sec. S15 we provide some references to literature about Bayesian variable selection.

### S1. From discrete-time branching process to diffusion

We provide an overview of the population genetics based approach we take to formulating a likelihood that connects observed count data to unobserved selection coefficients. For additional details (in somewhat different notation) please refer to Lee et al. (2022).

#### S1.1. Negative Binomial branching process

A natural model of viral infection is to suppose that an infected individual at time step $t$ infects a stochastic number of individuals in the next time step, where the number of infected individuals is governed by a Negative Binomial distribution with mean $R$ and variance $R + R^2/k$, where $k$ is the dispersion parameter. In the limit that $k \to \infty$ we recover a Poisson distribution with mean and variance both equal to $R$. In the opposite 'super-spreading' limit, $k \to 0$, the Negative Binomial distribution has a (potentially very large) variance that scales like $k^{-1}$. We now suppose that there are $n$ infected individuals at time step $t$. Under this assumption the total number of infected individuals at time step $t+1$ is governed by a Negative Binomial distribution with mean $nR$ and variance given by $(nR) + (nR)^2/(nk) = nR + nR^2/k$.

To make the model more realistic we suppose there are $V$ distinct viral variants in circulation. Each viral variant $v$ has an effective reproduction number given by $R_v = R_0(1 + \Delta R_v)$ where $R_0$ corresponds to the wild type. At the current time step $t$ there are $n_v(t)$ individuals infected with variant $v$. Given our assumptions, the number of infected individuals with variant $v$ at the next time step is governed by the distribution

$$n_v(t+1) \sim \text{NegativeBinomial}(\text{mean} = n_v(t)R_v, \text{dispersion} = n_v(t)k) \tag{1}$$

where we have assumed that $k$ is the same for all $V$ variants and disregard the possibility of mutation.

#### S1.2. Diffusion in variant frequency space

Following (Lee et al., 2022) we now transform from case counts $n_v(t)$ to variant frequencies $z_v(t)$:

$$z_v(t) \equiv \frac{n_v(t)}{\sum_{v'} n_{v'}(t)} \tag{2}$$

In the limit of large population size, i.e. the diffusion limit $n \gg 1$, and under the assumption that $|\Delta R_v| \ll 1$, Lee et al. (2022) compute approximate first and second moments for the variant frequencies $z_v(t)$ under the dynamics in Eqn. 1. These moments can in turn be used to define an (approximately) equivalent diffusion process for continuous-valued variant frequencies $z_v(t) \in [0, 1]$. In particular one finds that $\mathbf{z}(t)$ is approximately distributed like a Multivariate-Normal random variable as

$$\mathbf{z}(t+1) \sim \mathcal{N}(\mathbf{z}(t) + \tilde{\mathbf{d}}(t), \nu^{-1}\tilde{\mathbf{\Lambda}}(t)) \tag{3}$$

where $\tilde{\mathbf{d}}(t) \in \mathbb{R}^V$ is the $V$-dimensional drift

$$\tilde{d}_v(t) = z_v(t)\left(\Delta R_v - \sum_w \Delta R_w z_w(t)\right) \tag{4}$$

and $\tilde{\mathbf{\Lambda}}(t)$ is the $V \times V$ (unscaled) diffusion matrix given by

$$\tilde{\Lambda}_{v,w}(t) = \begin{cases} z_v(t)(1 - z_v(t)) & \text{if } v = w \\ -z_v(t)z_w(t) & \text{if } v \neq w \end{cases} \tag{5}$$

and $\nu$ is the effective population size given by

$$\nu \equiv \left(\tfrac{1}{R_0} + \tfrac{1}{k}\right)^{-1} n = \tfrac{kR_0}{k+R_0} n \tag{6}$$

Eqn. 3 has a simple and intuitive form with the properties we would expect. For example, the drift $\tilde{\mathbf{d}}(t)$ equals zero and thus the mean of $\mathbf{z}(t+1)$ is equal to $\mathbf{z}(t)$ if all variants have the same reproduction number (i.e. $\Delta R_v = 0$ for all $v$). Similarly, the variant $v$ with the largest $\Delta R_v$ will tend to see its prevalence increase, i.e. $\mathbb{E}[z_v(t+1) - z_v(t)] > 0$. For fixed $n$ the effective population size $\nu$ decreases as $k$ decreases, i.e. as the dynamics become increasingly characterized by super-spreading and are thus increasingly stochastic. In particular large populations experience approximately deterministic dynamics dominated by drift, while small populations experience considerably more stochastic dynamics in which the drift is more difficult to discern.

### S1.3. Diffusion in allele frequency space

Following (Lee et al., 2022), it is convenient to transform to a representation based on allele frequencies so that we can model allele-level selection. Suppose the virus has $A$ alleles with time-dependent allele frequencies $0 \leqslant x_a(t) \leqslant 1$ for $a = 1, \ldots, A$, i.e. $x_a(t)$ is the fraction of infected individuals at time $t$ infected by a virus that carries allele $a$. Denote pairwise allele frequencies as $x_{ab}(t)$, i.e. $x_{ab}(t)$ is the fraction of infected individuals at time $t$ infected by a virus that carries both allele $a$ and allele $b$. For convenience we define $x_{aa}(t) \equiv x_a(t)$. Further suppose that each variant $v$ is given as a genotype vector $\mathbf{g}_v \in \{0,1\}^A$, i.e. $g_{v,a}$ encodes whether variant $v$ carries allele $a$ ($g_{v,a} = 1$) or not ($g_{v,a} = 0$). We also assume that the variant-level differential reproduction number $\Delta R_v$ is governed by a linear additive model in allele-space:

$$\Delta R_v = \sum_a g_{v,a} \beta_a \tag{7}$$

where $\beta_a \in \mathbb{R}$ is an allele-level selection coefficient with positive (respectively, negative) coefficients corresponding to increased (decreased) viral fitness.

With these ingredients in hand and under the linear assumption in Eqn. 7 we can now specify the dynamics in allele frequency space, where the dynamics in Eqn. 3 are transformed to

$$\mathbf{x}(t+1) \sim \mathcal{N}(\mathbf{x}(t) + \mathbf{d}(t), \nu^{-1}\mathbf{\Lambda}(t)) \tag{8}$$

where $\mathbf{d}(t) \in \mathbb{R}^A$ is the $A$-dimensional drift,[1] given by

$$\mathbf{d}(t) = \mathbf{\Lambda}(t)\boldsymbol{\beta} \qquad \Longleftrightarrow \qquad d_a(t) = x_a(t)(1 - x_a(t))\beta_a + \sum_{b \neq a} \left(x_{ab}(t) - x_a(t)x_b(t)\right)\beta_b \tag{9}$$

and $\nu$ is as in Eqn. 6. Here $\mathbf{\Lambda}(t)$, analog to the diffusion matrix in Eqn. 5, is given by the $A \times A$ matrix

$$\Lambda_{ab}(t) = x_{ab}(t) - x_a(t)x_b(t) \tag{10}$$

Note that whenever alleles $a$ and $b$ exhibit non-trivial correlation, i.e. whenever $x_{ab}(t) \neq x_a(t)x_b(t)$, $d_a(t)$ depends on $\beta_b$ (in addition to $\beta_a$). In other words, a neutral allele can be 'dragged along' (i.e. exhibit apparent drift) by a non-neutral allele via hitchhiking. This effect is sometimes called *genetic draft*.

It is convenient to introduce incremental allele frequency changes

$$\mathbf{y}(t) \equiv \mathbf{x}(t+1) - \mathbf{x}(t) \tag{11}$$

so that Eqn. 8 becomes

$$\mathbf{y}(t) \sim \mathcal{N}(\mathbf{d}(t), \nu^{-1}\mathbf{\Lambda}(t)) \tag{12}$$

Eqn. 12, together with the definitions in Eqn. 9-11, forms the basis of everything that follows.

### S1.4. Relationship to PyR$_0$

An alternative approach to inferring allele-level selection effects is the PyR$_0$ model described in (Obermeyer et al., 2022). At its core this hierarchical Bayesian model relies on the following multivariate logistic growth ansatz

$$z_v(t) = \frac{e^{\Delta R_v t}}{\sum_{v'} e^{\Delta R_{v'} t}} \tag{13}$$

where $\mathbf{z}(t)$ are variant frequences like in Eqn. 3 and $\Delta R_v$ are variant-level differential growth rates. In direct analogy to the approach adopted in this work as well as Lee et al. (2022) PyR$_0$ proceeds to regress $\Delta R_v$ against allele-level features (cf. Eqn. 7). How is the PyR$_0$ ansatz in Eqn. 13 related to the diffusion-based dynamics in Eqn. 3? Eqn. 3 can be viewed as a specification of a stochastic differential equation in which there is deterministic drift $\tilde{\mathbf{d}}(t)$ as well as stochastic diffusion controlled by $\tilde{\mathbf{\Lambda}}(t)$. In the

[1] Note that Sec. S1.2 and Sec. S1.4 are the only sections in the supplement where we consider diffusion in variant-frequency space. In particular elsewhere drift vectors and diffusion matrices always correspond to diffusion in allele-frequency space.

infinite population limit, $\nu \to \infty$, the dynamics are deterministic and reduce to an ordinary differential equation. It is easy to check that Eqn. 13 is a solution to precisely that differential equation, namely

$$\frac{d}{dt}\mathbf{z}(t) = \tilde{\mathbf{d}}(t) \tag{14}$$

It would thus appear that the PyR$_0$ likelihood can be viewed as the deterministic counterpart of the diffusion-based likelihood in Eqn. 3. However, this is not quite correct, since PyR$_0$ uses the ansatz in Eqn. 13 to parameterize the probabilities of a $V$-dimensional Multinomial distribution. That is, PyR$_0$ describes an infinite population undergoing deterministic dynamics, with only a finite number of viral sequences observed at each time point. As a consequence[2] the noise structure of the PyR$_0$ Multinomial likelihood mirrors that of Eqn. 3, with the difference that the Multinomial likelihood implicitly assumes a fixed population size of $n$, where $n$ is the total number of observed viral sequences at a given time step. Importantly, it does not allow for the reduction in effective population size that results from super-spreading (small $k$, see Eqn. 6) or a finite sampling rate (see Sec. S8).

As we will see in our analysis of SARS-CoV-2 data, the effective population size in real data is quite modest, even in relatively well-sampled regions. Thus the stochastic component of the dynamics is expected to play an important role and, we would argue, should be accounted for explicitly in the model using a data-driven estimate of the effective population size. The ability to do so is one of the main benefits of using a diffusion-based likelihood.

### S2. Maximum a posteriori inference (MAP)

The simplest model that utilizes the diffusion-based likelihood in Eqn. 12 is formulated as follows (we refer the reader to Lee et al. (2022) for additional discussion). First we place a Multivariate-Normal prior on the $A$-dimensional vector of selection coefficients $\boldsymbol{\beta}$

$$p(\boldsymbol{\beta}|\tau) = \mathcal{N}(\boldsymbol{\beta}|\mathbf{0}, \tau^{-1}\mathbb{1}_A) \tag{15}$$

where $\tau > 0$ is the prior precision on the coefficients and $\mathbb{1}_A$ is the $A \times A$ identity matrix. For observed incremental allele frequency changes $\mathbf{y}(t)$ for $t = 1, ..., T-1$ the likelihood is given by

$$p(\mathbf{y}_{1:T-1}|\boldsymbol{\beta}, \nu) = \prod_{t=1}^{T-1} \mathcal{N}(\mathbf{d}(t|\boldsymbol{\beta}), \nu^{-1}\boldsymbol{\Lambda}(t)) \tag{16}$$

where we have assumed that $\nu$ is constant across time. Since $\boldsymbol{\beta}$ appears linearly in the drift $\mathbf{d}(t|\boldsymbol{\beta})$ (see Eqn. 9) and the prior is Multivariate-Normal, the corresponding posterior distribution, which is given by

$$p(\boldsymbol{\beta}|\mathbf{y}_{1:T-1}, \nu, \tau) = \frac{\mathcal{N}(\boldsymbol{\beta}|\mathbf{0}, \tau^{-1}\mathbb{1}_A) \prod_{t=1}^{T-1} \mathcal{N}(\mathbf{d}(t|\boldsymbol{\beta}), \nu^{-1}\boldsymbol{\Lambda}(t))}{\int d\boldsymbol{\beta}' \mathcal{N}(\boldsymbol{\beta}'|\mathbf{0}, \tau^{-1}\mathbb{1}_A) \prod_{t=1}^{T-1} \mathcal{N}(\mathbf{d}(t|\boldsymbol{\beta}'), \nu^{-1}\boldsymbol{\Lambda}(t))} \tag{17}$$

can be computed in closed form and is itself a Multivariate-Normal distribution. In particular the posterior mean, which in this case is also the maximum a posteriori (MAP) estimate, is given by

$$\boldsymbol{\beta}^{\mathrm{MAP}} \equiv \arg\max_{\boldsymbol{\beta}} p(\boldsymbol{\beta}|\tau) p(\mathbf{y}_{1:T-1}|\boldsymbol{\beta}, \nu) \tag{18}$$

$$= \mathbb{E}_{p(\boldsymbol{\beta}|\mathbf{y}_{1:T-1}, \nu, \tau)} [\boldsymbol{\beta}] \tag{19}$$

$$= \left( \sum_{t=1}^{T-1} \boldsymbol{\Lambda}(t) + \frac{\tau}{\nu}\mathbb{1}_A \right)^{-1} \sum_{t=1}^{T-1} \mathbf{y}(t) \tag{20}$$

$$= \left( \sum_{t=1}^{T-1} \boldsymbol{\Lambda}(t) + \frac{\tau}{\nu}\mathbb{1}_A \right)^{-1} \big(\mathbf{x}(T) - \mathbf{x}(1)\big) \tag{21}$$

We note that the effective regularization parameter for MAP is given by the ratio between the prior precision $\tau$ and the effective population size $\nu$:

$$\gamma_{\mathrm{reg}} \equiv \frac{\tau}{\nu} \tag{22}$$

An attractive property of this estimator is that it can be computed in $\mathcal{O}(A^3)$ time and is thus quite fast on modern hardware, at least for $A$ up to $A \sim 10^4 - 10^5$. An unattractive property of this estimator is that it can perform poorly in the high-dimensional case, $A \gg 1$, since we expect most alleles to be neutral, but $\beta_a^{\mathrm{MAP}}$ will generally be non-zero for all $a$.

[2] Note that a Multinomial distribution with $n$ trials and event probabilities $p_v$ for $v = 1, ..., V$ has variance $np_v(1-p_v)$ and covariance $-np_v p_w$ (for $v \neq w$).

### S3. Bayesian Viral Allele Selection

We expect most alleles to be neutral ($\beta_a = 0$) or nearly neutral ($\beta_a \approx 0$) and we would like to explicitly include this assumption in our model. To do so we utilize the modeling motif of Bayesian Variable Selection (Chipman et al., 2001). In more detail we consider the following space of models:

$$\begin{aligned} &[\text{inclusion variables}] && \gamma_a \sim \text{Bernoulli}(h) && \text{for} && a = 1, \ldots, A \\ &[\text{selection coefficients}] && \boldsymbol{\beta}_\gamma \sim \mathcal{N}(0, \tau^{-1}\mathbb{1}_{|\gamma|}) && && \\ &[\text{allele frequency changes}] && \mathbf{y}(t) \sim \mathcal{N}(\mathbf{d}(t|\boldsymbol{\beta}_\gamma), \nu^{-1}\boldsymbol{\Lambda}(t)) && \text{for} && t = 1, \ldots, T-1 \end{aligned} \tag{23}$$

Here each Bernoulli latent variable $\gamma_a \in \{0, 1\}$ controls whether the $a^{\text{th}}$ coefficient $\beta_a$ is included ($\gamma_a = 1$) or excluded ($\gamma_a = 0$) from the model; in other words it controls whether the $a^{\text{th}}$ allele is neutral ($\gamma_a = 0$) or not ($\gamma_a = 1$). In the following we use $\gamma$ to refer to the full $A$-dimensional vector $(\gamma_1, \ldots, \gamma_A)$. The hyperparameter $h \in (0, 1)$ controls the overall level of sparsity; in particular $hA$ is the expected number of non-neutral alleles a priori.[3] The $|\gamma|$ coefficients $\boldsymbol{\beta}_\gamma \in \mathbb{R}^{|\gamma|}$ are governed by a Normal prior with precision $\tau$ where $\tau > 0$ is a fixed hyperparameter. Here $|\gamma| \in \{0, 1, \ldots, A\}$ denotes the total number of non-neutral alleles in a given model. Finally we note that the drift $d(t|\boldsymbol{\beta}_\gamma)_a$ is given by $\boldsymbol{\Lambda}(t)_{a\gamma}\boldsymbol{\beta}_\gamma$, i.e. all alleles $b$ not included in the model (alleles with $\gamma_b = 0$) can be understood as having null coefficients $\beta_b = 0$. *In the following we drop the $\gamma$ subscript on $\boldsymbol{\beta}_\gamma$ to simplify the notation.*

In addition to inducing sparsity, an attractive feature of the model in Eqn. 23 is that—because it is formulated as a model selection problem—it explicitly reasons about whether each allele is neutral or not. In particular this model allows us to compute the *Posterior Inclusion Probability* or PIP, an interpretable score that satisfies $0 \leqslant \text{PIP} \leqslant 1$. The PIP is defined as

$$\text{PIP(a)} \equiv \text{p}(\gamma_\text{a} = 1|\mathbf{y}_{1:\text{T}-1}, \nu) \tag{24}$$

i.e. $\text{PIP}(a)$ is the posterior probability that allele $a$ is included in the model. This quantity should be contrasted to $h$ in Eqn. 23, which is the *a priori* inclusion probability. Alleles that have large PIPs are good candidates for being causally linked to viral fitness.

### S4. MCMC Inference

In this section we describe our approach to inference for the model in Sec. S3. Before we do so it is worth emphasizing that this is a difficult inference problem. There are two basic reasons for this. First, this is a transdimensional inference problem defined on a mixed discrete/continuous latent space. In particular the dimension of $\boldsymbol{\beta}$ depends on $|\gamma|$. Second, the size of the model space is astronomically large whenever there are more than a few dozen alleles. Indeed for $A$ alleles the total number of distinct models, namely $2^A$, exceeds the estimated number of atoms in the known universe ($\sim 10^{80}$) for $A \gtrsim 266$. However, the model also exhibits conjugacy structure that we can exploit. In particular, as we discuss next, $\boldsymbol{\beta}$ can be integrated out analytically for any given $\gamma$.

#### S4.1. Integrating out $\beta$

Recall from Eqn. 9 that the drift is given by $\mathbf{d}(t) = \boldsymbol{\Lambda}(t)\boldsymbol{\beta}$ so that each term in the likelihood (see Eqn. 12) can be written as

$$(\mathbf{y}(t) - \mathbf{d}(t))^{\text{T}}\boldsymbol{\Lambda}(t)^{-1}(\mathbf{y}(t) - \mathbf{d}(t)) = \tag{25}$$

$$\mathbf{d}(t)^{\text{T}}\boldsymbol{\Lambda}(t)^{-1}\mathbf{d}(t) - 2\mathbf{d}(t)^{\text{T}}\boldsymbol{\Lambda}(t)^{-1}\mathbf{y}(t) = \boldsymbol{\beta}^{\text{T}}\boldsymbol{\Lambda}(t)\boldsymbol{\beta} - 2\boldsymbol{\beta}^{\text{T}}\mathbf{y}(t) \tag{26}$$

where we have ignored the pre-factor of $-\frac{\nu}{2}$ and dropped terms that do not depend on $\boldsymbol{\beta}$. After re-introducing $-\frac{\nu}{2}$, each term in our likelihood contains the following quadratic form:

$$-\tfrac{\nu}{2}\left(\boldsymbol{\beta}^{\text{T}}\boldsymbol{\Lambda}(t)\boldsymbol{\beta} - 2\boldsymbol{\beta}^{\text{T}}\mathbf{y}(t)\right) \tag{27}$$

With this formula in hand we can compute the log marginal likelihood in closed form for any particular $\gamma$; up to irrelevant constants we have:

$$\log p(\mathbf{y}_{1:T-1}|\gamma) = \tfrac{1}{2}\bar{\mathbf{y}}_\gamma^{\nu\text{T}}\left(\bar{\boldsymbol{\Lambda}}^\nu_{\gamma\gamma} + \tau\mathbb{1}_{|\gamma|}\right)^{-1}\bar{\mathbf{y}}^\nu_\gamma - \tfrac{1}{2}\log\det\left(\bar{\boldsymbol{\Lambda}}^\nu_{\gamma\gamma} + \tau\mathbb{1}_{|\gamma|}\right) + \tfrac{1}{2}\log\det\left(\tau\mathbb{1}_{|\gamma|}\right) \tag{28}$$

where

$$\bar{\mathbf{y}}^\nu \equiv \nu\sum_{t=1}^{T-1}\mathbf{y}(t) = \nu(\mathbf{x}(T) - \mathbf{x}(1)) \qquad \bar{\boldsymbol{\Lambda}}^\nu \equiv \nu\sum_{t=1}^{T-1}\boldsymbol{\Lambda}(t) \tag{29}$$

Eqn. 28 can be computed efficiently using basic linear algebra tricks described in e.g. Zanella and Roberts (2019) and Jankowiak (2021). It is also important to note that $\bar{\mathbf{y}}^\nu$ and $\bar{\boldsymbol{\Lambda}}^\nu$ only need to be computed once before we run MCMC inference. In particular this means that MCMC iteration cost does not depend on $T$, since the summation over $t$ in Eqn. 29 can be done in pre-processing.

Crucially, since $\boldsymbol{\beta}$ can be integrated out analytically, our inference problem reduces to an inference problem over $\gamma$-space. This is convenient because $\gamma \in \{0, 1\}^A$ has fixed dimensionality and so we no longer need to concern ourselves with transdimensional inference.

[3] For more discussion see Sec. S9 below.

#### S4.2. MCMC

The resulting inference problem is still difficult, since the posterior over $\gamma$ is defined on a space of size $2^A$. Gibbs sampling offers a simple approach to inference for this model. However, Gibbs sampling mixes excruciatingly slowly unless $A$ is very small. To obtain an efficient algorithm we utilize a recently introduced MCMC inference algorithm, Tempered Gibbs Sampling, described in Zanella and Roberts (2019).[4] Here we give a very high-level description of this algorithm and refer the reader to Zanella and Roberts (2019) for more details.

We emphasize two main features of this algorithm. The first feature is that the MCMC algorithm targets an auxiliary posterior distribution that is tempered w.r.t. our original target distribution. Samples from the original target distribution are then obtained by reweighting, i.e. importance sampling. Crucially the algorithm is designed in such a way that tempering dramatically improves mixing while the variance of the importance weights remains moderate. Second, the algorithm makes extensive use of the fact that Eqn. 28 can be computed in closed form. In particular it makes use of the fact that, for any given value of $\gamma$, it is relatively cheap to compute the quantity $p(\gamma_a|\gamma_{-a}, \mathbf{y}_{1:T-1})$ for all $a$ in parallel. Here $\gamma_{-a}$ denotes all the entries of $\gamma$ except for $a$. This information is then used by the algorithm to make greedy moves in the immediate neighborhood of $\gamma$. Consequently the algorithm is able to quickly and efficiently explore regions of $\gamma$ space with high posterior probability.

#### S4.3. Computational complexity

The computational complexity of a single MCMC iteration is given by $\mathcal{O}(|\gamma|^3 + |\gamma|^2 A)$, where $|\gamma|$ is the number of non-neutral alleles at a given iteration and $A$ is the total number of alleles. If we assume that the sparsity assumption is valid so that $|\gamma| \ll A$ the computational complexity simplifies to $\mathcal{O}(|\gamma|^2 A)$ and is dominated by the cost of computing $p(\gamma_a|\gamma_{-a}, \mathbf{y}_{1:T-1})$ for $a = 1, ..., A$. In practice we find that this is quite fast on a mid-grade GPU. For example, for $A = 2904$ and using a NVIDIA Tesla T4 GPU it takes about 45 minutes to generate half a million MCMC samples, which is about 200 samples per second.

### S5. Multiple spatial regions

Above we have assumed that observed allele frequencies $\mathbf{x}(t)$ are from a single spatial region. The generalization to multiple regions is straightforward. First we add a $r = 1, ..., N_R$ subscript to the likelihood:

$$\mathbf{y}_r(t) \sim \mathcal{N}(\mathbf{d}_r(t), \nu_r^{-1}\mathbf{\Lambda}_r(t)) \quad \text{for} \quad r = 1, ..., N_R \quad \text{and} \quad t = 1, ..., T-1 \tag{30}$$

Depending on the strategy used to estimate the effective population size (see Sec. S6) each $\nu_r$ in Eqn. 30 is either distinct or is instead a single global value. The equation for the marginal log likelihood, Eqn. 28, is unchanged except we now write Eqn. 29 as

$$\bar{\mathbf{y}}^\nu \equiv \sum_{r=1}^{N_R} \nu_r \sum_{t=1}^{T-1} \mathbf{y}_r(t) \qquad\qquad \bar{\mathbf{\Lambda}}^\nu \equiv \sum_{r=1}^{N_R} \nu_r \sum_{t=1}^{T-1} \mathbf{\Lambda}_r(t) \tag{31}$$

This equation makes it evident that our effective population size estimates play an important role, since they control the relative contribution of different regions to the region-summed totals in $\bar{\mathbf{y}}^\nu$ and $\bar{\mathbf{\Lambda}}^\nu$. In particular using a per-region $\hat{\nu}$ estimator has the consequence that regions with larger effective population sizes contribute more to the likelihood than regions with small effective population sizes. This is of course as expected, since regions with smaller effective population sizes are more stochastic and thus contribute less certain information about the drift. We discuss our strategy for estimating $\nu$ in the next section.

### S6. Inferring the effective population size $\nu$

The likelihood in Eqn. 12 depends on the effective population size $\nu$, a quantity that we do not know a priori and need to estimate from data. By assumption the increment $\mathbf{y}(t)$ is distributed as

$$\mathbf{y}(t) \sim \mathcal{N}(\mathbf{d}(t), \nu^{-1}\mathbf{\Lambda}(t)) \tag{32}$$

Under this assumption

$$\mathbb{E}\left[\mathbf{y}(t)^{\mathrm{T}}\mathbf{y}(t)\right] = \mathbf{d}(t)^{\mathrm{T}}\mathbf{d}(t) + \nu^{-1}\mathrm{Tr}\mathbf{\Lambda}(\mathrm{t}) \tag{33}$$

If we assume that the drift term is subdominant so that

$$\mathbb{E}\left[\mathbf{y}(t)^{\mathrm{T}}\mathbf{y}(t)\right] \approx \nu^{-1}\mathrm{Tr}\mathbf{\Lambda}(\mathrm{t}) \tag{34}$$

this leads to the estimator

$$\hat{\nu} = \frac{\mathrm{Tr}\mathbf{\Lambda}(\mathrm{t})}{\mathbb{E}\left[\mathbf{y}(t)^{\mathrm{T}}\mathbf{y}(t)\right]} \tag{35}$$

We note that, since $\mathbf{d}(t)^{\mathrm{T}}\mathbf{d}(t) \geqslant 0$, we might expect $\hat{\nu}$ to be an underestimate of $\nu$.

[4] In particular we use the `wTGS` variant and make use of Rao-Blackwellized PIP estimators.

Thus for a particular region we can estimate the effective population size as

$$\hat{\nu} = \tfrac{1}{T-1} \sum_{t=1}^{T-1} \frac{\mathrm{Tr}\boldsymbol{\Lambda}(\mathrm{t})}{\mathbf{y}(t)^{\mathrm{T}}\mathbf{y}(t)} \tag{36}$$

where we have averaged Eqn. 35 over $T-1$ time steps. We can also partially correct for the bias introduced by the (a priori unknown and non-zero) drift by instead using a centered estimator:

$$\hat{\nu} = \tfrac{1}{T-1} \sum_{t=1}^{T-1} \frac{\mathrm{Tr}\boldsymbol{\Lambda}(\mathrm{t})}{\left(\mathbf{y}(t) - \tfrac{1}{T-1}\bar{\mathbf{y}}\right)^{\mathrm{T}}\left(\mathbf{y}(t) - \tfrac{1}{T-1}\bar{\mathbf{y}}\right)} \tag{37}$$

with

$$\bar{\mathbf{y}} \equiv \sum_{t=1}^{T-1} \mathbf{y}(t) = \mathbf{x}(T) - \mathbf{x}(1) \tag{38}$$

denoting time-integrated allele increments. We find that both estimators in Eqn. 36 and Eqn. 37 work well in simulation, but for simplicity we use Eqn. 36 throughout this work. Indeed it is also possible to construct $\hat{\nu}$ estimators that are more time-local in nature and can thus accommodate population sizes that change over time, but for simplicity we prefer to use the hyperparameter-free estimator in Eqn. 36, as we expect this choice to be more robust. More complex schemes are certainly possible, but it is likely that they would require additional sources of data beyond (partially observed) genomic surveillance data if they are to be reliable (e.g. case count data).

The discussion above assumes a single spatial region. How can we accommodate multiple regions? We adopt two possible schemes. In the *global* scheme we compute $\hat{\nu}$ within each region using Eqn. 36 and then compute a single global effective population size $\hat{\nu}$ via a mean or median over all regions. In the *regional* scheme we simply compute $\hat{\nu}$ within each region using Eqn. 36. Since we expect the latter scheme to lack robustness when confronted with the complexities of real-world SARS-CoV-2 data, we only make comparisons to the regional scheme in simulation.

### S7. Incorporating time-dependent vaccination rates

Suppose we know the vaccination rate $0 \leqslant \phi_r(t) \leqslant 1$ for a given region $r$. We would like to incorporate this information into our modeling. In particular we would like to allow for vaccination-dependent selection. To do this we write the drift in region $r$ as

$$d_{r,a}(t) = x_{r,a}(t)(1 - x_{r,a}(t))\left(\beta_a + \phi_r(t)\alpha_a\right) + \sum_{b \neq a} \left(x_{r,ab}(t) - x_{r,a}(t)x_{r,b}(t)\right)\left(\beta_b + \phi_r(t)\alpha_b\right) \tag{39}$$

where $\boldsymbol{\alpha} \in \mathbb{R}^A$ is a second group of selection coefficients whose strength is modulated by the time- and region-local vaccination rate $\phi_r(t)$. In particular $\boldsymbol{\alpha}$ only has a non-negligible effect on viral dynamics when the vaccination rate has attained a non-negligible level. Disentangling the effects of $\boldsymbol{\beta}$ and $\boldsymbol{\alpha}$ is difficult a priori. However, our hope is that a Bayesian variable selection approach with robust MCMC inference should be up to the task provided we have enough data.

The particular form of Eqn. 39 can be motivated as follows. Eqn. 39 implies that for any given allele there are two selection coefficients: one for unvaccinated populations ($\beta$) and another for vaccinated populations ($\alpha + \beta$). These population-level effects can be connected back to the individual level. In particular take the $k \to \infty$ limit of the discrete time process in Sec. S1 so that secondary infections are controlled by a Poisson distribution. Suppose we are considering a variant with a single non-wild-type allele and there are $n$ infected individuals at the current time step. If we suppose that secondary infections result from contact between infected and non-infected individuals, and that an infected individual comes into contact with non-infected individuals who are vaccinationed/unvaccinated in the proportion $\phi$ to $1-\phi$, then the total number of secondary infections in the next time step is given by

$$n_{\mathrm{unvac}} + n_{\mathrm{vac}} \quad \text{with} \quad n_{\mathrm{unvac}} \sim \mathrm{Poisson}(R_0(1+\beta)(1-\phi)n) \quad \text{and} \quad n_{\mathrm{vac}} \sim \mathrm{Poisson}(R_0(1+\beta+\alpha)\phi n) \tag{40}$$

which is governed by a $\mathrm{Poisson}(R_0(1+\beta+\phi\alpha)n)$ distribution. Here $n_{\mathrm{unvac}}$ and $n_{\mathrm{vac}}$ are the number of secondary infections among unvaccinated and vaccinated individuals, respectively. Thus the effective selection coefficient is given by $\beta + \phi\alpha$, explaining the form of Eqn. 39.

The basic inference approach introduced in Sec. S4 is still applicable. All we need to do is suitably modify $\bar{\mathbf{y}}^\nu$ and $\bar{\boldsymbol{\Lambda}}^\nu$ in Eqn. 31. In particular we now have that $\bar{\mathbf{y}}^\nu \in \mathbb{R}^{2A}$

$$\bar{\mathbf{y}}^\nu \equiv \begin{pmatrix} \sum_{r=1}^{N_R} \nu_r \sum_{t=1}^{T-1} \mathbf{y}_r(t) \\ \sum_{r=1}^{N_R} \nu_r \sum_{t=1}^{T-1} \phi_r(t)\mathbf{y}_r(t) \end{pmatrix} \tag{41}$$

and $\bar{\boldsymbol{\Lambda}}^\nu \in \mathbb{R}^{2A \times 2A}$:

$$\bar{\boldsymbol{\Lambda}}^\nu \equiv \begin{pmatrix} \sum_{r=1}^{N_R} \nu_r \sum_{t=1}^{T-1} \boldsymbol{\Lambda}_r(t) & \sum_{r=1}^{N_R} \nu_r \sum_{t=1}^{T-1} \phi_r(t)\boldsymbol{\Lambda}_r(t) \\ \sum_{r=1}^{N_R} \nu_r \sum_{t=1}^{T-1} \phi_r(t)\boldsymbol{\Lambda}_r(t) & \sum_{r=1}^{N_R} \nu_r \sum_{t=1}^{T-1} \phi_r^2(t)\boldsymbol{\Lambda}_r(t) \end{pmatrix} \tag{42}$$

This change approximately doubles the cost of MCMC inference and quadruples memory requirements, but this regime is still well within reach of a mid-grade GPU even for $A \sim 10^4$.

### S8. Sampling rate

What effect does sampling—the fact that not all viral sequences are observed in real world datasets—have on our diffusion-based likelihood? We first give a somewhat high-level analysis that ignores some of the subtleties involved. Suppose that each infected individual has their viral genome sequenced with probability $0 < \rho < 1$. Due to sampling, a variant $v$ that is circulating in $n_v$ individuals will be observed $\tilde{n}_v \sim \text{Binomial}(n_v, \rho)$ times. This distribution has mean $\rho n_v$ and variance $\rho(1-\rho)n_v$. For $\rho \ll 1$ and large $n_v$ we have that $\tilde{n}_v$ is very nearly governed by the Poisson distribution $\text{Poisson}(\rho \mathrm{n_v})$, which has mean and variance both equal to $\rho n_v$. Note, however, that the Poisson distribution is just the $k \to \infty$ limit of the Negative Binomial distribution, which in turn is the basic ingredient of the discrete time process that underlies our diffusion-based likelihood (see Eqn. 1). For this reason the combined action of one step of viral infection and sampling rate on the allele frequencies $x_a(t)$ can be described by the equation

$$\mathbf{x}(t+1) \sim \mathbf{x}(t) + \mathbf{d}(t) + \boldsymbol{\epsilon} + \boldsymbol{\epsilon}' \tag{43}$$

with

$$\boldsymbol{\epsilon} \sim \mathcal{N}(\mathbf{0}, \nu^{-1}\boldsymbol{\Lambda}(t)) \qquad \boldsymbol{\epsilon}' \sim \mathcal{N}(\mathbf{0}, \nu'^{-1}\boldsymbol{\Lambda}(t))) \tag{44}$$

where

$$\nu' \equiv \rho n \tag{45}$$

is the effective population size corresponding to variability due to sampling and where $\boldsymbol{\epsilon}$ and $\boldsymbol{\epsilon}'$ encode variability in infection and sampling, respectively. Since, however, both sources of variability are zero mean normally distributed noise in the diffusion limit, they can easily be combined:

$$\mathbf{x}(t+1) \sim \mathbf{x}(t) + \mathbf{d}(t) + \boldsymbol{\epsilon}'' \qquad \text{with} \qquad \boldsymbol{\epsilon}'' \sim \mathcal{N}(\mathbf{0}, \nu''^{-1}\boldsymbol{\Lambda}(t))) \tag{46}$$

where

$$\nu''^{-1} = \nu^{-1} + \nu'^{-1} \tag{47}$$

or in other words the effect of $\rho$ would appear to be to redefine the effective population size as

$$\nu \to \left(\tfrac{1}{R_0} + \tfrac{1}{k} + \tfrac{1}{\rho}\right)^{-1} n \tag{48}$$

As one would expect, for fixed $R_0$ and $k$ decreasing $\rho$ decreases the effective population size, since the variability due to sampling increases.

#### S8.1. Detailed analysis

While the simplified analysis above is already adequate for understanding the basic dynamics, it is necessary to do a more careful analysis to reveal some of the additional subtleties involved. In particular let us be careful about distinguishing between the unobserved allele frequencies $\tilde{x}_a(t)$ (corresponding to all infected individuals) and the observed allele frequencies $x_a(t)$ (corresponding to infected individuals whose genomes are sequenced). We can write

$$\begin{aligned}
&\text{[infection]} && \tilde{\mathbf{x}}(t+1) \sim \mathcal{N}(\tilde{\mathbf{x}}(t) + \tilde{\mathbf{d}}(t), \nu^{-1}\tilde{\boldsymbol{\Lambda}}(t)) && (49)\\
&\text{[sampling]} && \mathbf{x}(t) \sim \mathcal{N}(\tilde{\mathbf{x}}(t), \nu'^{-1}\tilde{\boldsymbol{\Lambda}}(t)) && (50)\\
&\text{[sampling]} && \mathbf{x}(t+1) \sim \mathcal{N}(\tilde{\mathbf{x}}(t+1), \nu'^{-1}\tilde{\boldsymbol{\Lambda}}(t+1)) && (51)
\end{aligned}$$

where $\tilde{\mathbf{d}}(t)$ and $\tilde{\boldsymbol{\Lambda}}(t)$ are defined in terms of $\tilde{\mathbf{x}}(t)$ and $\tilde{\boldsymbol{\Lambda}}(t+1)$ is defined in terms of $\tilde{\mathbf{x}}(t+1)$, and where we have assumed that $\rho$ and thus $\nu'$ is constant across time. Ultimately we are interested in how $\mathbf{x}(t+1)$ depends on $\mathbf{x}(t)$, since these are the quantities we observe. To make progress in analyzing Eqn. 49-51 we first make the replacements $\tilde{\mathbf{d}}(t) \to \mathbf{d}(t)$, $\tilde{\boldsymbol{\Lambda}}(t) \to \boldsymbol{\Lambda}(t)$ and $\tilde{\boldsymbol{\Lambda}}(t+1) \to \boldsymbol{\Lambda}(t+1)$, which we expect to be approximately valid in the diffusion limit $n \to \infty$.[5] With this substitution we find that $\mathbf{x}(t+1)$ is distributed as

$$\mathbf{x}(t+1) \sim \mathcal{N}(\mathbf{x}(t) + \mathbf{d}(t), \nu^{-1}\boldsymbol{\Lambda}(t) + \nu'^{-1}\boldsymbol{\Lambda}(t) + \nu'^{-1}\boldsymbol{\Lambda}(t+1)) \tag{52}$$

If we now further assume that the drift is sufficiently small so that $\boldsymbol{\Lambda}(t) \approx \boldsymbol{\Lambda}(t+1)$ we find that

$$\mathbf{x}(t+1) \sim \mathcal{N}(\mathbf{x}(t) + \mathbf{d}(t), \nu''^{-1}\boldsymbol{\Lambda}(t)) \tag{53}$$

where

$$\nu'' = \left(\tfrac{1}{R_0} + \tfrac{1}{k} + \tfrac{2}{\rho}\right)^{-1} n \tag{54}$$

so that Eqn. 48 was almost correct, except that—since we need to 'denoise' twice, i.e. once for time step $t$ and once for time step $t+1$—the $\rho^{-1}$ term appears with an additional factor of two. It is easy to check numerically that the approximation in Eqn. 53 is a reasonably good one provided that $n$ is large, $\rho$ is small, and the drift is subdominant.

[5] By Eqn. 50-51 $\mathbf{x}(t) \to \tilde{\mathbf{x}}(t)$ and $\mathbf{x}(t+1) \to \tilde{\mathbf{x}}(t+1)$ as $n \to \infty$.

Eqn. 53 is reassuring because it means that sampling induces noise that is congruent with the noise induced by the Negative Binomial dynamics that underlies the discrete time process in Sec. S1. In particular this means that we do not need to separately estimate e.g. $k$ and $\rho$, since it suffices to estimate the effective population size in Eqn. 54. This is a significant simplification and is a key component in the viability of our overall approach.

### S9. Prior inclusion probability

An important hyperparameter in our modeling approach is the prior inclusion probability $h$ in Eqn. 23. In an equivalent parameterization we can instead consider the quantity

$$S \equiv \mathbb{E}\left[\sum_a \gamma_a\right] = hA \tag{55}$$

where $S$ is the expected number of non-neutral alleles a priori and the expectation in Eqn. 55 is with respect to the prior in Eqn. 23.

We consider two approaches to setting $h$ (and thus $S$). The first is to set $h$ by hand utilizing any a priori knowledge we may have of the expected number of non-neutral alleles. While a priori knowledge of $h$ may be imprecise, this need not be particularly troubling: as we show in simulations in the main text results are not expected to be overly sensitive to the precise value of $h$. Indeed, it is worth emphasizing that $h$ is the *prior* inclusion probability. Sufficient evidence in the likelihood can always overwhelm the prior and the posterior inclusion probability (the PIP) can differ significantly from $h$.

The second approach is to place a prior on $h$ (and thus implicitly on $S$). A natural choice is to place a Beta prior on $h$

$$h \sim \mathrm{Beta}(\alpha_h, \beta_h) \tag{56}$$

where $\alpha_h > 0$ and $\beta_h > 0$ are hyperparameters that control the Beta prior. This prior assumption has the following properties. The prior mean of $h$ is given by

$$\mathbb{E}[h] = \frac{\alpha_h}{\alpha_h + \beta_h} \tag{57}$$

For $\beta_h \gg \alpha_h$ and $\beta_h \gg 1$ this mean is approximately equal to $\alpha_h/\beta_h$ and we further have that the variance of $h$ is approximately given by

$$\mathbb{V}[h] \approx \frac{\alpha_h}{\beta_h^2} \tag{58}$$

Consequently in this regime the ratio of the square root variance to the mean—which is a useful measure of the width of the prior distribution—is approximately given by the formula

$$\frac{(\mathbb{V}[h])^{\frac{1}{2}}}{\mathbb{E}[h]} \approx \frac{1}{\sqrt{\alpha_h}} \tag{59}$$

These formulae are useful for choosing $\alpha_h$ and $\beta_h$. For example, if we have $A = 1000$ alleles and we expect that $S$ is about 10 and likely between 3 and 20 we might choose $\alpha_h = 1$ and $\beta_h = 100$, which concentrates about 90% of the prior probability mass between $S = 0.5$ and $S = 30$.

Once we have chosen $\alpha_h$ and $\beta_h$ we must still infer $h$. In the absence of tempering this is easy to do since the posterior conditional distribution over $h$ is just another Beta distribution due to conjugacy. Unfortunately, the tempering introduced by the MCMC scheme we use spoils this relationship (Zanella and Roberts, 2019). Nevertheless this conjugacy can easily be restored by introducing an additional auxiliary latent state in the MCMC algorithm as explained in detail in Jankowiak (2021). Thus in practice inferring $h$ is straightforward.

### S10. Alternative modeling and inference strategies

In Sec. S1 we described how a diffusion-based likelihood can be used to model genomic surveillance data. In Sec. S2 we described the simplest inference strategy that utilizes such a likelihood, namely MAP inference. In Sec. S3 we described our own diffusion-based approach: Bayesian Viral Allele Selection. This method differs from MAP in two notable ways: i) it uses a different prior; and ii) it uses a different inference algorithm (MCMC instead of maximum a posteriori inference). In this section we describe a few alternative diffusion-based modeling and inference strategies.[6]

[6] To lighten the notation we assume a single spatial region.

#### S10.1. Laplace prior

We first describe the simplest modification of MAP that can account for the expected sparsity of non-neutral alleles. In this approach we place a Laplace prior on $\boldsymbol{\beta}$

$$p(\boldsymbol{\beta}|\sigma^{\text{Laplace}}) = \frac{1}{2\sigma^{\text{Laplace}}} \exp\left(-\frac{||\boldsymbol{\beta}||_1}{\sigma^{\text{Laplace}}}\right) \tag{60}$$

where $||\boldsymbol{\beta}||_1$ is the $L^1$ norm of $\boldsymbol{\beta}$ and $\sigma^{\text{Laplace}} > 0$ is a hyperparameter that controls the expected level of sparsity. We then define the maximum a posteriori estimate under this Laplace prior:[7]

$$\boldsymbol{\beta}^{\text{Laplace}} \equiv \underset{\boldsymbol{\beta}}{\arg\max}\, p(\boldsymbol{\beta}|\sigma^{\text{Laplace}}) p(\mathbf{y}_{1:T-1}|\boldsymbol{\beta}, \nu) \tag{61}$$

In order to compute $\boldsymbol{\beta}^{\text{Laplace}}$ we need to rely on optimization methods. In our experiments we use the Adam optimizer (Kingma and Ba, 2014) and implement inference using the Pyro probabilistic programming framework (Bingham et al., 2019). In particular we use the Adam optimizer with an initial learning rate of $10^{-2}$ that exponentially decays to $10^{-4}$ over the course of $10^4$ optimization steps.[8] Solving this optimization problem is $\mathcal{O}(A^3 + A^2 T_{\text{opt}})$ where $T_{\text{opt}}$ is the total number of optimization steps and the cubic cost in $A$ results from computing the Cholesky factorization of $\boldsymbol{\Lambda}^{\nu}$.

Another possible approach would be to consider the exact same model as in Eqn. 61 but to perform variational inference, for example mean-field variational inference utilizing a product of Normal distributions as the variational distribution over $\boldsymbol{\beta}$. We would expect this approach to suffer from one of the main issues that plagues the approach in Eqn. 61: the hyperparameter $\sigma^{\text{Laplace}}$ is difficult to set, but a well-chosen value of $\sigma^{\text{Laplace}}$ is crucial for good performance.

#### S10.2. Horseshoe prior

Another approach to inducing shrinkage in $\boldsymbol{\beta}$ would be to use a hierarchical shrinkage prior like the Horseshoe (Carvalho et al., 2009). However, in our experience MCMC algorithms for Horseshoe-based regression can mix quite poorly in the high-dimensional setting. In addition, since the Horseshoe prior is a continuous shrinkage prior, it does not provide an interpretable allele-level score like the PIP provided by a Bayesian Variable Selection approach (see Eqn. 24). Moreover, MCMC inference in this context is likely to be expensive, since $\boldsymbol{\Lambda}^{\nu}$ does not have low-rank structure that can be exploited for computational speed-ups.

#### S10.3. Discussion

Having now discussed a few possible alternatives to BVAS, some of the favorable characteristics of BVAS come into sharper focus. First, a fully Bayesian approach that systematically integrates over different hypotheses about which alleles are neutral and which are not contributes to reduced sensitivity to prior hyperparameters. Second, the sparsity assumption has notable computational benefits with the consequence that BVAS admits efficient MCMC inference with $\mathcal{O}(|\gamma|^2 A)$ cost per iteration (e.g. compare to the quadratic dependence on $A$ in the Laplace method). Third, BVAS offers interpretable posterior inclusion probabilities. Fourth, the MCMC inference algorithm itself does not include any difficult-to-set hyperparameters, in contrast for example to the Laplace method above, which may require tweaking optimization hyperparameters for optimal performance.

### S11. Note on Statistical Analysis

All statistical analysis in this work is done using standard Bayesian methods. In particular unless noted otherwise we report 95% credible intervals based on the (approximate) posterior distribution. Since all our models jointly consider all data—as opposed to univariate association tests—no corrections for multiple hypothesis testing are necessary.

### S12. Limitations

We provide an extended discussion of some of the limitations of BVAS not touched upon in the main text. Due to limitations of our pre-processing pipeline that we inherit from UShER, we do not consider insertions and deletions. If there are insertions or deletions that have non-negligible selection effects, the exclusion of these alleles from our analysis may lead to inferences that incorrectly implicate other alleles that are linked to these insertions/deletions. These considerations are also applicable to our epistasis analysis, which may be particularly sensitive to any such missing allelic information. More broadly, the fidelity of the epistasis analysis may also be impacted by the exclusion of interaction effects outside the RBD.

Our method cannot correct for any issues in the raw input data (e.g. sequencing errors) and cannot correct for lineage-dependent sampling bias (e.g. sequencing only in case of S-gene target failure). Likewise biased inferences can result if, e.g., disease severity varies across variants and disease severity influences the likelihood that an individual's viral genome is sequenced. Our diffusion-based likelihood cannot account for differential fitness effects that depend on unobserved confounders like population age structure. In addition we assume that individuals are infected with a single SARS-CoV-2 variant, an assumption that can be violated (Chan et al., 2022).

[7] For the sake of precision we should probably refer to the approach in Sec. S2 as MAP-Gaussian and the approach described here as MAP-Laplace. However, for brevity we instead refer to these methods as MAP and Laplace, respectively. One crucial difference between these two methods is that the former method admits a closed form solution, whereas the latter method requires iterative optimization techniques.

[8] The hyperparameters that control the momentum are $(\beta_1, \beta_2) = (0.5, 0.99)$, which is appropriate for this non-stochastic regime.

### S13. Simulations

#### S13.1. Simulation details (extended)

Unless otherwise noted, the parameters and distributional assumptions of all simulation-based experiments are as follows. We consider $V = 100$ viral variants and a true selection coefficient vector $\boldsymbol{\beta}^*$ with 10 non-zero coefficients:

$$\boldsymbol{\beta}^* \equiv (0.01, 0.02, 0.04, 0.06, 0.08, -0.01, -0.02, -0.04, -0.06, -0.08, -0.10, 0, 0, 0, ..., 0) \tag{62}$$

In other words there are exactly 10 non-neutral alleles, 5 of which have positive effects on viral fitness and 5 of which have negative effects on viral fitness. We sample each element of each genotype vector $\mathbf{g}_v$ i.i.d. from a Bernoulli distribution with mean $\frac{1}{5}$, i.e. $g_{v,a} \sim \text{Bernoulli}(\frac{1}{5})$ for $v = 1, ..., V$ and $a = 1, ..., A$ where $A$ is the total number of alleles. This choice implies that typical variants have about two non-neutral alleles. We set $R_0 = 1$ so that typical reproduction numbers for variants $v$ range between $R_v \sim 0.9$ and $R_v \sim 1.1$. In each simulation we consider a given number of $R$ regions and $T = 26$ time steps. This latter choice corresponds to about one year of data assuming time bins of 14 days.

The initial number of infected individuals at time $t = 1$ within each region is drawn from a Negative Binomial distribution with mean $10^4$ and dispersion parameter[9] 10. Within each region these infected individuals are then apportioned to each of the $V$ variants using a Multinomial distribution with uniform probabilities $V^{-1}$. Thus the mean case count of each variant $v$ within each region at time $t = 1$ is given by $10^4/V = 100$. Case counts for $t = 2, ..., T$ are then determined by the stochastic dynamics in Eqn. 1 with $k = 0.1$. This value of $k$ is chosen since it consistent with estimates of the SARS-CoV-2 dispersion parameter (Lau et al., 2020; Endo et al., 2020; Bi et al., 2020; Miller et al., 2020). These raw counts are then subjected to Binomial sampling with mean $\rho = 0.01$, representing a sampling rate of 1%, i.e. the viral sequences of 99% of cases are not observed. Note that this implies that the expected number of *observed* individuals with variant $v$ within each region at the initial time step is 1. Thus our parameter choices result in simulated data that is highly stochastic and that constitute a regime in which we expect that recovering the true selection coefficients $\boldsymbol{\beta}^*$ is quite challenging. Unless noted otherwise, we generate 20 datasets per condition.

We make these choices because they result in simulated data that exhibit some of the characteristics of our SARS-CoV-2 data. In particular, typical estimated effective population sizes $\hat{\nu}$ range from about 25 to about 140 with a mean of about 75. Indeed from Eqn. 54 we would expect that

$$\hat{\nu} \approx \left(\tfrac{1}{1} + \tfrac{1}{0.1} + \tfrac{2}{0.01}\right)^{-1} \times 10^4 \approx \tfrac{0.01}{2} \times 10^4 \approx 50 \tag{63}$$

which is indeed what we observe, see Figure S1. Note the fact that most regions have an effective population size somewhat larger than 50 is due to the fact that in most regions the number of infected individuals increases throughout the course of the pandemic.

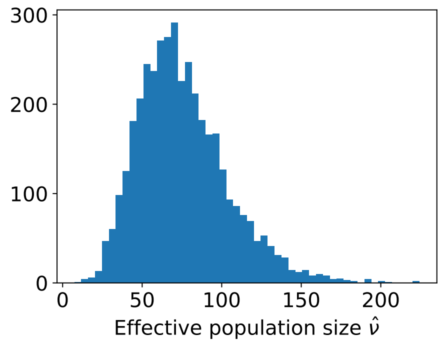


**Fig. S1.** We depict the distribution of estimated effective population sizes $\hat{\nu}$ for the simulated data described in Sec. S13.1. The mean and standard deviation are 75.6 and 28.1, respectively. Note that the underlying true population size is about $10^4$.

#### S13.2. $\hat{\nu}$ estimator performance

We investigate the accuracy of the effective population size estimators described in Sec. S6. Our simulated data are as in Sec. S13.1 except that we limit the duration of the pandemic to $T = 10$ time steps to limit the exponential growth/decay within any given region so that the true population size is approximately constant across time within any given region. We run the simulator for $N_R = 10^5$ regions. We then use Eqn. 54 to compute the 'true' effective population size for each region, where we average Eqn. 54 across $T$ time steps (since the true number of infected individuals fluctuates from time step to time step). Finally we use our basic $\hat{\nu}$ estimator, Eqn. 36, to estimate the effective population size for each region.

In Figure S2 we depict histograms of ratios of the estimated $\hat{\nu}$ to the true effective population size $\nu_{\text{true}}$ for two scenarios. In the first scenario (left) we simulate data with $\boldsymbol{\beta} = \boldsymbol{\beta}^*$ as in Eqn. 62, with the result that drift is relatively moderate. In the second scenario (right) we simulate data with $\boldsymbol{\beta} = 5\boldsymbol{\beta}^*$, with the result that drift is much more pronounced. We find reasonably good agreement with a relative accuracy of about 10-40% depending on the scenario.

[9] The variance is thus $\sim 3164^2$.

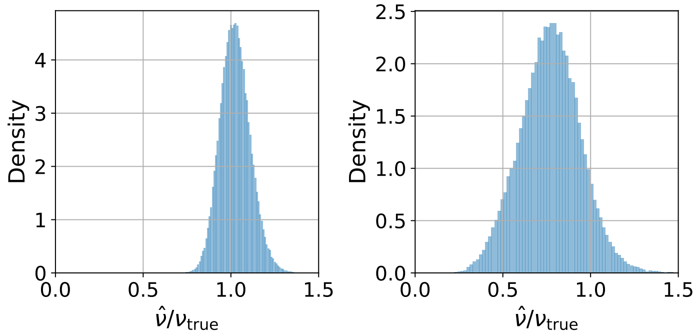


**Fig. S2.** We evaluate the performance of our basic $\hat{\nu}$ estimator, Eqn. 36. We compare a scenario with smaller drift (left) and larger drift (right). See Sec. S13.2 for details. The ratio is approximately normally distributed with a mean of 1.02 and standard deviation of 0.09 (left) and a mean of 0.77 and standard deviation of 0.17 (right). Typical reproduction numbers $R_v$ range from about 0.9 to 1.1 (left) and about 0.5 to 1.5 (right).

### S13.3. Method Comparison

We provide some additional information regarding the method comparison in the main text. We use the following hyperparameter settings. For MAP we test all values of $\tau$ ranging from $2^{-24}$ to $2^{24}$ in powers of 2 and choose $\tau = 2^{11} = 2048$, since this value gives the best results across the board. In other words we are using the ground truth to choose $\tau$, something that we obviously cannot do when analyzing real data. Since the effective regularization parameter for MAP is given by $\gamma_{\mathrm{reg}} = \tau/\nu$ (see Eqn. 22) and $\nu \approx 75$ for our simulated data this corresponds to a value of $\gamma \approx 30$, which is in the range of values suggested by Lee et al. (2022).[10] For BVAS we use $\tau = 100$ and $(\alpha_h, \beta_h) = (1/16, A/32)$ except for experiments where we explicitly vary the prior inclusion probability $h$.[11] Unless specified otherwise these BVAS settings are used in all simulation experiments. For Laplace we use a regularization scale of $\sigma^{\mathrm{Laplace}} = 0.01$. For $\mathrm{PyR}_0$ we use the default settings in Obermeyer et al. (2022). Unless specified otherwise, we use the global $\hat{\nu}$ estimator described in Sec. S6.[12] We note that we tried using z-scores to rank $\mathrm{PyR}_0$ hits but found essentially identical results. We also note that the hit rate can be understood as the precision that results if the top 10 alleles are declared hits. We run BVAS for 5500 iterations, discarding the first 500 samples for burn-in. We use $T_{\mathrm{opt}} = 10^4$ optimization steps when running Laplace.

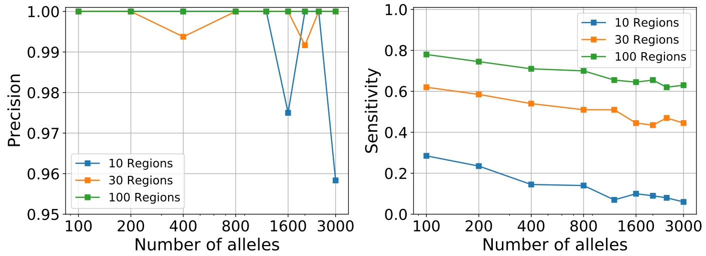


**Fig. S3.** We depict the precision and sensitivity performance of BVAS using the simulated data described in Sec. S13.1. For this purpose we define hits as all alleles with PIPs above a threshold of 0.1. Precision is the number of true positives divided by the total number of hits. Sensitivity is the fraction of causal alleles that are identified as hits.

### S13.4. Comparing global vs. regional $\hat{\nu}$ strategies

In the main text we investigated the effect of modulating the overall scale of $\nu$. It is also important to investigate whether we should prefer the global or regional $\hat{\nu}$ estimators introduced in Sec. S6. As can be seen in Figure S6, the two strategies perform nearly identically, at least on our simulated data. This suggests that the more 'ambitious' regional estimator *could* perform well in real data, but it also suggests that it may be entirely sufficient to use the (presumably) more robust global estimator. For this reason we only use the global estimator in our actual analysis. By virtue of this choice all regions in our analyses contribute equally to the likelihood.

[10] We found that it is only for $A = 100$ where there is a strong preference for a smaller value of $\tau$ like $\tau = 2^9$; for larger $A$ there is a uniform preference for larger values of $\tau$.

[11] This corresponds to approximately 2 expected non-neutral alleles a priori and encodes substantial a priori uncertainty about $h$. See Sec. S9 for discussion. Note that we use a prior over $h$ instead of specifying $h$ as a fixed hyperparameter to avoid leveraging the fact that we know that the true number of non-neutral alleles is given by 10.

[12] This is only relevant for the diffusion-based methods. The particular value of $\nu$ used is most relevant for BVAS and Laplace and is somewhat less important for MAP, which only depends on $\nu$ through the quantity $\gamma_{\mathrm{reg}} = \tau/\nu$.

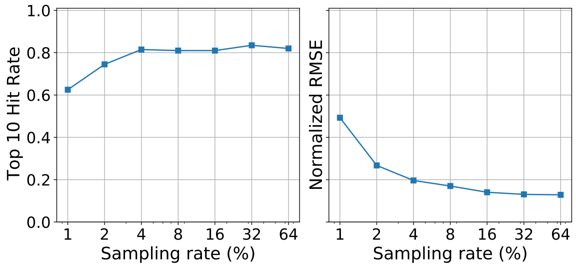


**Fig. S4.** We depict how the hit rate (left) and root mean squared error (RMSE; right) vary as a function of the sampling rate. We consider $A = 2000$ alleles and $N_R = 30$ regions.

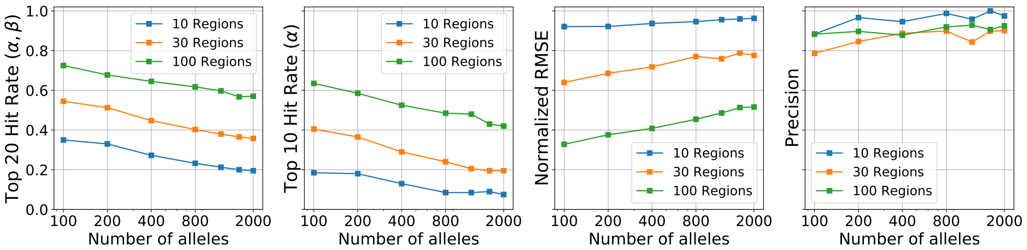


**Fig. S5.** We explore BVAS performance in a scenario with vaccination-dependent selection coefficients. Precision computations assume a PIP cutoff of 0.1. Note that we only count a hit as correct if it is correctly identified as $\phi$-dependent or $\phi$-independent.

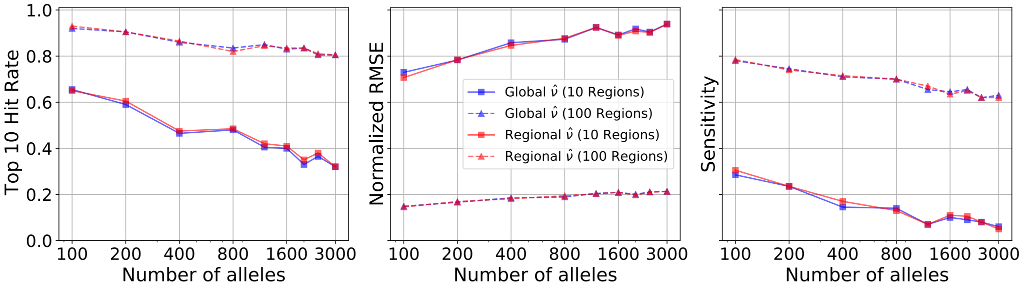


**Fig. S6.** We compare two different strategies for estimating the effective population size $\nu$ on simulated data. See Sec. S13.4 for details.

#### S13.5. Spike-in experiment

SARS-CoV-2 data are prepared as in Sec. S14.1. Then for each variant $v$ in the data the genotype $\mathbf{g}_v$ is extended by 200 fake spike-in alleles, where we sample the genotype of each spike-in allele i.i.d. from a Bernoulli distribution with mean 0.01.

#### S13.6. Including vaccination-dependent effects

Our simulation is as in Sec. S13.1 except we now assume 20 non-zero effects, 10 of which are vaccination-dependent and 10 of which are not (we assume these two sets of ten are disjoint and that each set has the same effect sizes as the non-zero effects in Eqn. 62). We assume that $\phi_r(t)$ starts at zero everywhere (i.e. $\phi_r(t=1)=0$), increases linearly over time, and that the final value of $\phi_r(t)$ in each region is given by $\phi_r(T) \sim \mathrm{Uniform}(\frac{1}{2}, 1)$.

### S14. SARS-CoV-2 Analysis

#### S14.1. Data Pre-processing

The initial steps of our data pre-processing pipeline follow the procedures described in Obermeyer et al. (2022) and make use of the open source code at `https://github.com/broadinstitute/pyro-cov`. We give a short summary of the relevant parts of this pipeline and refer the reader to Obermeyer et al. (2022) for additional details.

Our initial data ingest consists of 8,575,902 samples downloaded from GISAID (Elbe and Buckland-Merrett, 2017) on April 18$^{\text{th}}$, 2022. Each sample record includes metadata for time of collection, geographic location, and PANGO lineage annotation (Rambaut et al., 2020) as well as genetic sequence data. We discard records without complete metadata. Sequences whose alignment quality is not reported as “good” are discarded. Time intervals are binned into 14-day segments, which results in a total of 62 time intervals, i.e. a bit more than two years of data. Note that we choose a multiple of 7 days to minimize weekly seasonality effects (e.g. closure of testing facilities on weekends).

#### *S14.1.1. Lineage Clustering*

Since the likelihood that underlies BVAS is defined in allele frequency space, it does not explicitly rely on notions of variant or lineage. However, the pre-processing pipeline in Obermeyer et al. (2022) makes use of a dynamic partitioning of genetic samples into clusters. One potential advantage of this approach is that it can remove sequencing artifacts. Like Obermeyer et al. (2022) our default analysis makes use of $L = 3000$ clusters derived from the 1694 PANGO lineages present in our dataset. That PANGO lineages are not sufficiently fine can be seen in Figure S7, which shows that some PANGO lineages (notably B.1.1) consist of several sublineages with widely varying fitnesses. This recapitulates the findings in Obermeyer et al. (2022). These $L = 3000$ clusters are created as follows. The starting point is a 9,293,486 node phylogeny of all GISAID samples maintained by Angie Hinrichs (McBroome et al., 2021). This phylogeny was created using UShER (Turakhia et al., 2021) and excludes private mutations, masks difficult-to-sequence regions, disregards deletions, and parsimoniously imputes missing sequence data. To coarse grain the 9.3 million node phylogenetic tree down to $L = 3000$ clusters we iteratively collapsed parent-child edges by greedily minimizing a notion of distance defined between pairs of mutation-annotated trees. Empirically Obermeyer et al. (2022) found that this heuristic clustering produces trees that are approximately balanced (both in terms of cluster size and cluster-cluster edit distance). Note that this clustering procedure removes some of the genetic diversity in the raw data. The total number of alleles retained at the end of clustering is $A = 2975$. In Figure S20 we investigate the sensitivity of our analysis to the choice of $L$ and find only modest differences.

#### *S14.1.2. Spatial Aggregation*

To accommodate the fact some spatial regions are sparsely sampled while other spatial regions are densely sampled, we dynamically aggregate regions to encourage *approximately* balanced sample totals per region. For example, in well-sampled countries like the US and Germany our spatial regions are at the state/province level (e.g. California and Bavaria) while subregions of less densely sampled countries are aggregated up to the country level (e.g. Turkey, Portugal, and Slovenia).

The result of the above pre-processing steps is a collection of region-specific time series of lineage counts (encoded as a $R \times T \times L$ count array) as well as lineage-level amino acid features (encoded as a $L \times A$ binary array with $A = 2975$).

#### *S14.1.3. Allele Frequency Space*

Running BVAS MCMC requires two basic ingredients (see Eqn. 28): $\bar{\mathbf{y}}^{\nu}$ and $\bar{\mathbf{\Lambda}}^{\nu}$. Since these allele frequency space quantities are unique to BVAS and the method in Lee et al. (2022) the rest of our pre-processing pipeline diverges from Obermeyer et al. (2022).

#### *S14.1.4. Diffusion Limit Filters*

BVAS relies on the diffusion limit, i.e. it assumes that the number of infected individuals is relatively large. For this reason we would like to exclude regions that are poorly sampled from our analysis.[13] With that caveat in mind we would like to include as many regions as possible so that the results of our analysis are not unduly influenced by the idiosyncracies of any particular region. To manage this trade-off we introduce two hyperparameters: $N_{\rm min}^{\rm tot}$ and $N_{\rm min}^{14}$. Regions where the total number of samples is less than $N_{\rm min}^{\rm tot}$ are not included in our analysis. By making $N_{\rm min}^{\rm tot}$ relatively large we exclude regions that have only been observed for a handful of time intervals (note that it is particularly difficult to estimate the effective population size of such regions). We must also decide which time intervals to include in our analysis. Even if the *total* number of samples in a given region is large the number of samples collected early on in the pandemic may be small, with the result that dynamics in that early phase of the pandemic will tend to be dominated by noise (i.e. diffusion). For each region we thus only include those 14-day time intervals for which the number of samples is greater than or equal to $N_{\rm min}^{14}$. Moreover, since our likelihood depends on allele frequency increments $\mathbf{y}(t)$ (see Eqn. 12) we choose the longest subsequence of contiguous time intervals where each time interval contains at least $N_{\rm min}^{14}$ samples.[14] In our default analysis, we choose relatively conservative values of these two hyperparameters: $N_{\rm min}^{\rm tot} = 10^4$ and $N_{\rm min}^{14} = 50$. In Sec. S14.2 we investigate the sensitivity of our estimates to $N_{\rm min}^{\rm tot}$ and $N_{\rm min}^{14}$. In Table S2 we list the 128 regions that enter into our default analysis with $N_{\rm min}^{\rm tot} = 10^4$ as well as the 67 regions that enter into an auxiliary analysis with $N_{\rm min}^{\rm tot} = 20 \times 10^3$.

#### *S14.1.5. Computing $\bar{\mathbf{y}}^{\nu}$ and $\bar{\mathbf{\Lambda}}^{\nu}$*

The next step is to compute $\bar{\mathbf{y}}^{\nu}$ and $\bar{\mathbf{\Lambda}}^{\nu}$. First we estimate the effective population size $\hat{\nu}$ for each region using the estimator in Eqn. 36. In our default global analysis we then compute the median $\hat{\nu}$ across all the regions included in the analysis (note that the number of regions included depends on $N_{\rm min}^{\rm tot}$) and scale the median value of $\hat{\nu}$ by a factor of 0.5. This choice is motivated by several concerns. The median effective population size $\hat{\nu}$ in our default analysis with 128 regions is 63.9. This value exceeds the estimated effective population size of 64 of the 128 regions, see Table S2. Moreover, half that value, namely 32.0, exceeds the estimated effective population size of only 9 of the 128 regions. Thus by making this choice we ensure that $\hat{\nu}$ is not grossly *over*estimated in any one region (the smallest estimated $\hat{\nu}$ is 20.0 in Fukuoka, Japan). Consequently, with this choice BVAS is not expected to confuse the inherent stochasticity of the least well sampled regions with signal.

---

13 In any case we do not expect to learn very much from regions with small effective population sizes, since the dynamics in these regions is dominated by noise. In this context see Figure S23 in Obermeyer et al. (2022), which shows that $\mathrm{PyR}_0$ results are largely driven by the most well-sampled regions.

14 This is especially important for accommodating vaccination-dependent effects as in Eqn. 42.

| | Growth Rate | | Growth Rate |
|---|---|---|---|
| **XN** | 7.328 ± 0.689 | **XM** | 6.934 ± 0.721 |
| **XT** | 7.273 ± 0.706 | **XK** | 6.736 ± 0.717 |
| **XR** | 7.249 ± 0.715 | **XS** | 6.293 ± 0.599 |
| **XL** | 7.206 ± 0.722 | **XP** | 6.216 ± 0.600 |
| **XG** | 7.197 ± 0.722 | **XD** | 5.848 ± 0.651 |
| **XQ** | 7.147 ± 0.715 | **XF** | 5.797 ± 0.575 |
| **XH** | 7.012 ± 0.718 | **XC** | 3.578 ± 0.487 |
| **XE** | 6.963 ± 0.722 | **XB** | 2.447 ± 0.406 |
| **XJ** | 6.962 ± 0.722 | **XA** | 2.268 ± 0.236 |

**Table S1.** We report the relative growth rates $R/R_A$ of recombinant SARS-CoV-2 lineages as estimated by BVAS. Uncertainty estimates denote 95% credible intervals.

To put this choice in a different light, we note that Bayesian inference has a long history of grappling with model mis-specification and robustness. One class of methods achieves robustness by raising the likelihood to a power, which is precisely the result of scaling the effective population size. Thus our choice of $\hat{\nu}$ can be understood as a way of achieving robustness against possible model mis-specification. In particular we do not expect our genomic surveillance data to exactly follow the discrete time branching process in Sec. S1 nor do we expect the reproduction number to be exactly linear in the genotype. For more discussion on these and related ideas see Bissiri et al. (2016).

With this choice of $\hat{\nu}$ we now compute $\bar{\mathbf{y}}^\nu$ and $\bar{\mathbf{\Lambda}}^\nu$ using the definitions in Eqn. 31 and using a shared value of the effective population size across all regions. Note that computing these quantities is $\mathcal{O}(N_R T)$, i.e. the computation scales with both the number of regions and the number of time intervals under consideration, but that $\bar{\mathbf{y}}^\nu$ and $\bar{\mathbf{\Lambda}}^\nu$ can be computed very efficiently in an online fashion.[15] Indeed computing $\bar{\mathbf{y}}^\nu$ and $\bar{\mathbf{\Lambda}}^\nu$ only takes a few seconds on a mid-grade GPU. Importantly, these $\mathcal{O}(N_R T)$ computations only need to be done once in pre-processing, since the BVAS MCMC algorithm operates directly on $\bar{\mathbf{y}}^\nu$ and $\bar{\mathbf{\Lambda}}^\nu$.

##### *S14.1.6. Vaccination rates*

Vaccination data are downloaded from OWID (Ritchie et al., 2020):

- `https://github.com/owid/covid-19-data/raw/master/public/data/vaccinations/vaccinations.csv`
- `https://github.com/owid/covid-19-data/raw/master/public/data/vaccinations/us_state_vaccinations.csv`

We use two definitions of vaccination status: `people_vaccinated_per_hundred` and `people_fully_vaccinated_per_hundred`. We use country-level data for all regions except for US states and England, Scotland, Wales. Any missing values are interpolated using linear interpolation. We consider the same 128 regions as in our default analysis, except we exclude Luxembourg, since it does not include complete vaccination information. Thus we consider 127 regions in total.

##### *S14.1.7. MCMC*

When running BVAS we do $51 \times 10^4$ MCMC iterations. We discard the first $10^4$ samples and retain the remaining $5 \times 10^5$ samples to compute posterior quantitities of interest like PIPs. All computations are done in 64-bit precision.

#### S14.2. Sensitivity Analysis

See Figure S13-S27 and the main text for discussion.

#### S14.3. Backtesting

We follow the same data pre-processing pipeline as in Sec. S14.1, although we note that for any given date fewer than 128 regions enter our analysis, since we are considering subsets of the data and the $N^{\rm tot}_{\rm min} = 10^4$ requirement is still in place. In practice a real-time surveillance program might choose to use looser filters (i.e. smaller $N^{\rm tot}_{\rm min}$ and $N^{14}_{\rm min}$).

#### S14.4. Comparison to MAP

In Figure S30-S35 we compare BVAS results to results obtained with MAP (Lee et al., 2022). Estimates for selection coefficients $\beta$ are *qualitatively* similar to BVAS results, although BVAS results are much sparser, with many coefficients nearly zero (Figure S30). Growth rate estimates are in good agreement if the MAP regularizer $\tau$ is set sufficiently low ($\tau = 2^8$ or $\tau = 2^{10}$), but otherwise MAP estimates appear to be over-regularized towards Wuhan A, see Figure S31-S32. Conversely, Manhattan plots for MAP estimates are dense unless $\tau \geqslant 2^{12}$ (Figure S34). Thus MAP is unable to simultaneously yield large growth estimates for fit lineages like Omicron *and* assign negligible effect sizes to most alleles. If we believe that most alleles should be neutral, this is a limitation of

[15] In particular we avoid instantiating large arrays whenever possible. For example, we do not compute $\bar{\mathbf{\Lambda}}^\nu$ separately for each region and then sum over regions in an outer loop; rather we accumulate a single $A \times A$ matrix as we go.

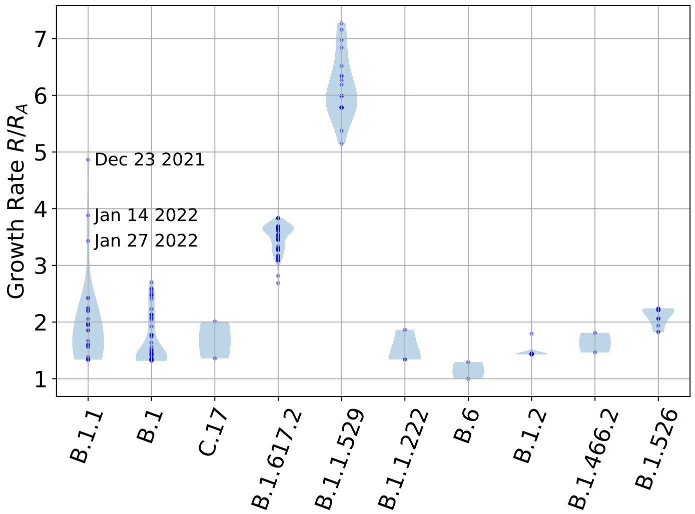


**Fig. S7.** We depict the 10 most diverse PANGO lineages as estimated by our model. The family of B.1.1 lineages is notably diverse, with some of the most recent sublineages—the dates of emergence for a few select lineages are annotated—exhibiting growth rates that exceed that of Delta and approach that of the least fit Omicron sublineages.

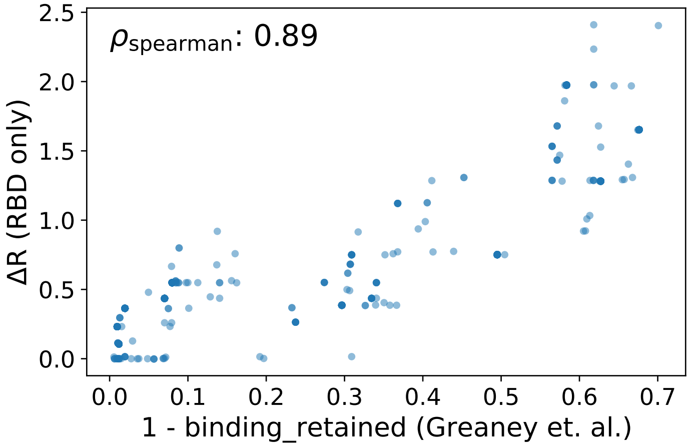


**Fig. S8.** For the 1729 SARS-CoV-2 clusters with at least one amino acid substitution in the RBD domain we compare: i) the BVAS prediction for the contribution to $\Delta R$ from RBD substitutions only; to ii) antibody binding computed using the antibody-escape calculator in Greaney et al. (2022). The escape calculator is based on deep mutational scanning data for 33 neutralizing antibodies elicited by SARS-CoV-2. BVAS predictions exhibit high (Spearman) correlation with predictions from Greaney et al. (2022). Compare to Figure 2B in Obermeyer et al. (2022).

MAP. More generally, MAP offers limited control over how much estimates should be regularized because it is controlled by a single hyperparameter $\tau$. This is in contrast to BVAS where regularization is controlled by both $S$ and $\tau$. Indeed BVAS uses a value of $\gamma_{\rm reg} \equiv \tau/\nu$ that is an order of magnitude smaller than MAP, depending on $S$ for additional regularization. The analysis in Figures S31-S34 indicates that this additional regularization is crucial. We also note that MAP uncertainty estimates are quite narrow (see e.g. Figure S31) which can ultimately be traced to the fact that MAP considers a single hypothesis about allele neutrality (namely that no alleles are neutral).

Note that our MAP analysis follows the exact same data pre-processing pipeline as for BVAS, see Sec. S14.1. Thus BVAS and MAP comparisons utilize the exact same likelihood, although we do not scale the $\hat{\nu}$ estimate in the case of MAP and instead use the median effective population size directly. Also note that in Figure S33 and Figure S35 we choose $\tau = 2^{12}$, which corresponds to a level of regularization that appears to achieve the best balance between over- and under-regularization—although as we have argued above this compromise appears to be suboptimal.

### S14.5. Comparison to PyR$_0$

We also compare BVAS results to results obtained with PyR$_0$, see Figure S36-S40. While PyR$_0$ hits cluster in similar parts of the SARS-CoV-2 genome as BVAS hits (Figure S36) and many of the top hits are common to both methods (Figure S37), the quantitative agreement between PyR$_0$ and BVAS is modest. For example, only 7 hits are common to the top 25 BVAS and

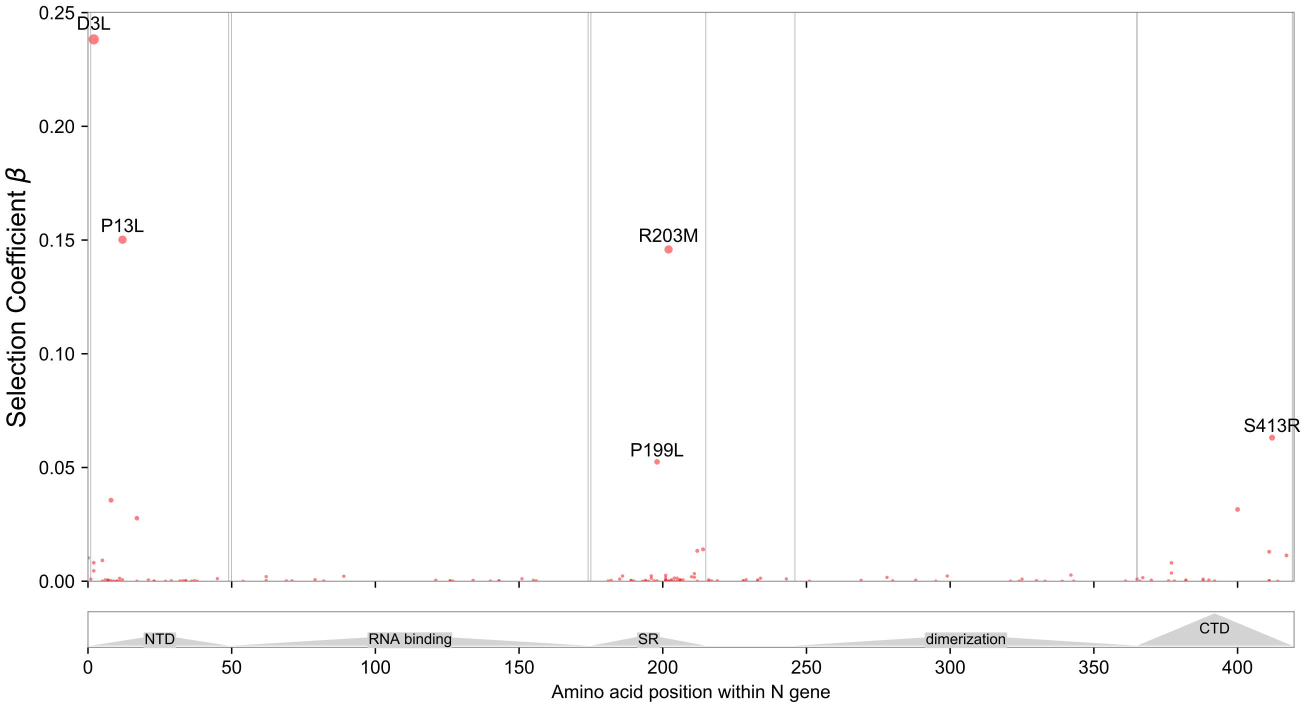


**Fig. S9.** View of the 419 amino acids of the nucleocapsid (N) protein domains, annotated by structure (Cubuk et al., 2021). Many top-scoring mutations occur in the serine–arginine rich region (SR), reported to be immunogenic by Chen et al. (2004).

top 25 $\text{PyR}_0$ hits, and the Pearson correlation for $\beta$ estimates is $R = 0.365$, see Figure S39. Moreover, $\text{PyR}_0$ estimates for the fittest Omicron lineages are substantially higher than BVAS estimates, by as much as $\sim 50\%$, see Figure S40. Additionally $\text{PyR}_0$ estimates, especially for allele-level quantities like selection coefficients, are implausibly small, orders of magnitude smaller than the corresponding BVAS uncertainties.

We note that, like our Laplace method (see Eqn. 60), $\text{PyR}_0$ has a hyperparameter $\sigma_3$ that controls the sparsity of the selection coefficients. The analysis in Obermeyer et al. (2022) uses $\sigma_3 = 5 \times 10^{-4}$. However, our analysis includes approximately $\sim 35\%$ more data than the analysis in Obermeyer et al. (2022), suggesting that a smaller value of $\sigma_3$ may be required. Consequently, after comparing inference results for several values, we settle on $\sigma_3 = 2 \times 10^{-4}$, although we note that other similar values are also likely defensible. This highlights one of the disadvantages of $\text{PyR}_0$: it depends on several hyperparameters that can be difficult to set in practice.

$\text{PyR}_0$ comparisons follow the data pre-processing pipeline described in Obermeyer et al. (2022). We use the default parameters listed in Obermeyer et al. (2022), with the difference that we decrease $\sigma_3$ (as discussed above). Note that unlike BVAS the $\text{PyR}_0$ analysis does not make use of filters like $N_{\text{min}}^{\text{tot}} = 10^4$, and thus considers more data from significantly more regions (1694 instead of 128).

### S14.6. Hitchhiking

As is well known, hitchhiking, i.e. apparent drift to due genetic linkage, complicates the inference of selection effects and makes it difficult to unambiguously disentangle driver and passenger mutations. Hitchhiking can involve both selective sweeps caused by the uptake of beneficial mutations (Smith and Haigh, 1974) as well as the elimination of neutral variants linked to deleterious mutations, i.e. background selection (Charlesworth et al., 1993). One of the advantages of the diffusion-based likelihood in Eqn. 12 is that covariance information from $\mathbf{\Lambda}(t)$ can help distinguish driver mutations from passenger mutations. To make this more explicit, we identify the 3 alleles that exhibit large uptakes in allele fractions while simultaneously exhibiting small PIPs below 0.01: N:R203K, ORF14:G50N, and N:G204R. As can be seen in Figure S41 these alleles have in fact peaked three times, having become very prevalent during the B.1.1, B.1.1.7 (Alpha), and BA.1 (Omicron) waves, while experiencing significant drops in interim periods when other lineages (e.g. B.1.177 and Delta) predominated. BVAS infers that these three alleles are passenger mutations whose rising and falling prevalences are best explained by selection effects due to other linked alleles. While it is difficult to validate whether this particular inference is correct, it is reassuring that all three alleles have similar PIPs, as they exhibit near perfect co-occurrence in the data.

### S14.7. Vaccination-dependent effects

We follow the same data pre-processing pipeline described in Sec. S14.1, noting that the dimension of $\mathbf{y}(t)$ doubles as we include vaccination-dependent effects. Note, however, that when we estimate the effective population size with Eqn. 36 we do so using allele frequency increments in the original $A$-dimensional space so that our $\hat{\nu}$ estimate remains the same.

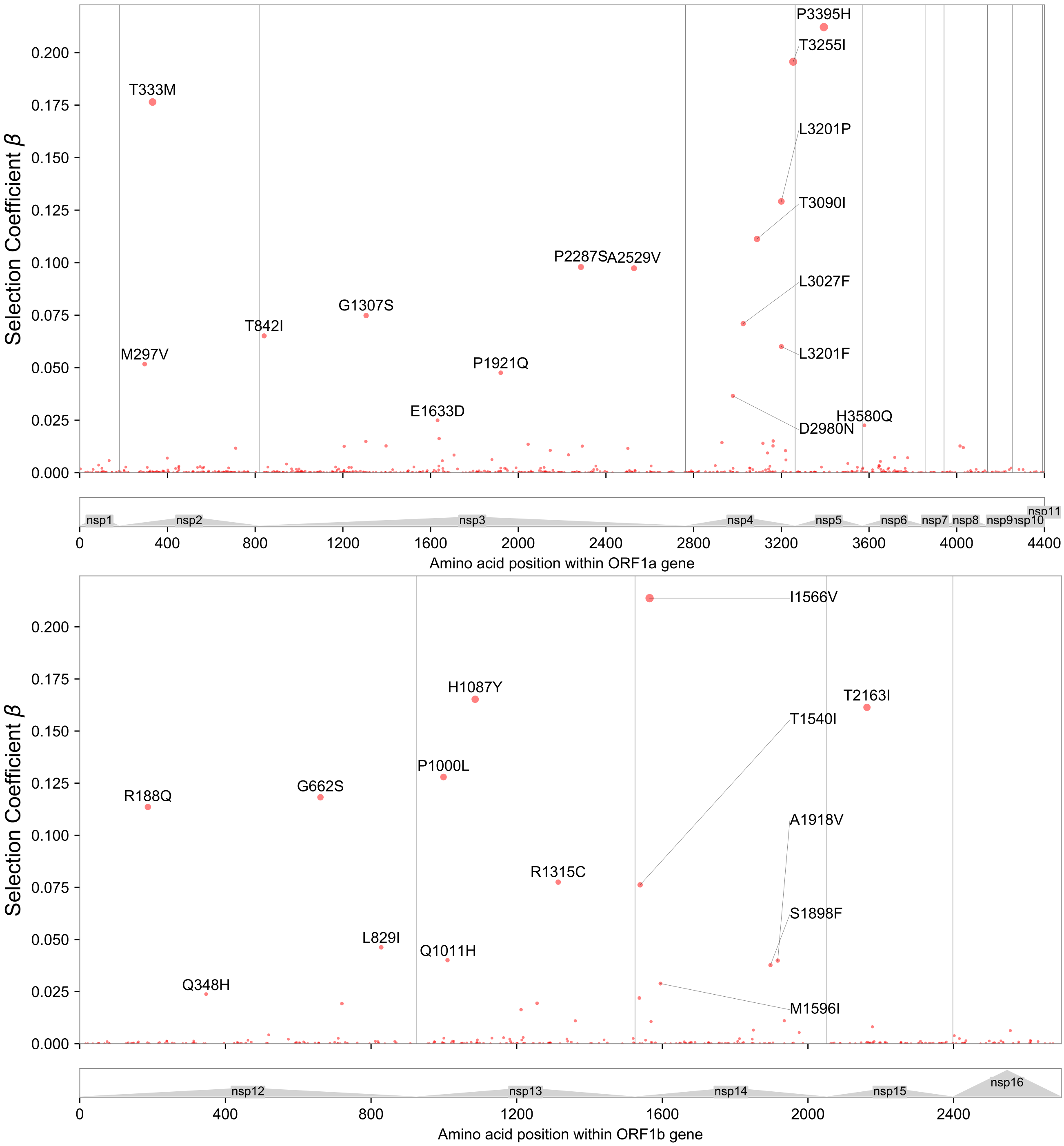


**Fig. S10.** We depict Manhattan plots zoomed-in on two regions of the SARS-CoV-2 genome. **(Top)** View of the ORF1a polyprotein, including 11 non-structural proteins (nsps). **(Bottom)** View of the ORF1b polyprotein, including nsp12-16; note the amino acid positions do not account for 9 additional residues at the N-terminus of nsp12 (RNA polymerase) resulting from the -1 ribosomal frameshift.

### S14.8. Epistasis

We report additional details pertaining to our epistasis analysis. We first note that the variant-space dynamics in Eqn. 3 are related to the allele-space dynamics in Eqn. 8 via a linear transformation specified by a genotype matrix. This still holds true when there are higher order interactions in the fitness ansatz in Eqn. 7. Consequently all the relevant formulae in Sec. S1 remain valid, with the difference that allele-space quantities for pairwise features must be re-interpreted appropriately. For example, entries $((a_1, a_2), (a_3, a_4))$ in the diffusion matrix $\mathbf{\Lambda}(t)$ in Eqn. 10 corresponding to pairwise features $(a_1, a_2)$ and $(a_3, a_4)$ encode *quartic* moments w.r.t. the underlying amino-acid-level alleles.

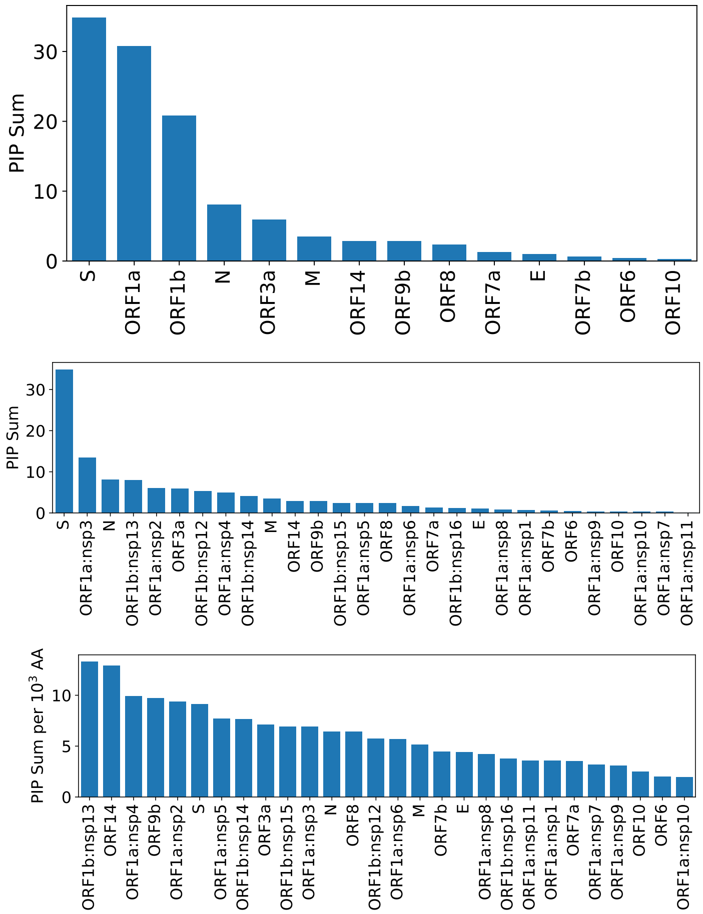


**Fig. S11.** As a measure of each gene's total contribution to selection we depict the summed PIP across all alleles in each gene in the SARS-CoV-2 genome (top). In the middle panel we break-up the two largest genes (ORF1a and ORF1b) into constituent NSPs (non-structural proteins). In the bottom panel we additionally normalize by the length of the gene (in units of 1000 amino acids).

Our epistasis analysis uses the same data and pre-processing as our default analysis. We use twice as many MCMC iterations as in our default analysis. For pairwise interaction effects we use an a priori inclusion probability $h_{\rm quad}$ that is one half of the value used for linear effects, i.e. $h_{\rm quad} = \frac{1}{2} h_{\rm linear} \approx 0.0084$. We verified that other values of $h_{\rm quad}$—in particular $h_{\rm quad} = \frac{1}{4} h_{\rm linear}$ and $h_{\rm quad} = h_{\rm linear}$—identified the same leading pairwise interaction effects (although with a corresponding shift in the overall PIP level for quadratic effects), with (N501Y, P681H) the leading hit in all three analyses. This remains true for sensitivity analyses in which $\hat{\nu}$ is modulated down by a factor of one half or the prior precision $\tau$ is halved.

As touched upon in the main text, we only consider quadratic interactions between pairs $(a_1, a_2)$ of non-synonymous amino acid substitutions that co-occur in at least 2 of our $L = 3000$ SARS-CoV-2 clusters. Moreover for each $(a_1, a_2)$ we require that there are at least two clusters that 'separate' the pair, i.e. clusters that carry the $a_1$ allele but not the $a_2$ allele or vice versa. After applying these filters there remain 1432 pairwise interaction effects for consideration, where the vast majority of pairwise interactions that do not pass our filters are excluded because of the co-occurrence requirement. Note that BVAS would not have any incentive to make use of these pairwise interactions if they were included in the analysis, since they would be unidentifiable in the context of their constituent linear effects (and as such they do not add any additional expressivity). See Figure S45 for a sensitivity analysis for (linear) effects common to the epistasis analysis and the default analysis without epistasis. Top hits from our epistasis analysis are reported in Table S4. The 'pairwise' posterior inclusion probabilities for these hits are depicted graphically in Figure S46.

In the main text we report that top-scoring interaction effects in our epistasis analysis are enriched for interactions between mutations that correspond to top-scoring (linear) effects in our default analysis. In particular 8 of the 10 amino acids in the top 8 quadratic hits in our epistasis analysis exhibit PIPs larger than 0.3 in our default analysis. Only A67V and E484A exhibit negligible PIPs in our default analysis but participate in a top 8 quadratic hit, namely the interaction between A67V and E484A.

The enrichment analysis is done as follows. For the 421 amino acid mutations $a$ in Spike under consideration, we assign a score given by $\sigma(a) = 1/\text{rank}$, where rank is the PIP rank of the corresponding linear effect in our default analysis (i.e. $\sigma(a) \in \{1/1, 1/2, 1/3, ...\}$). We then assign interaction effects between amino acids $a_1$ and $a_2$ a score given by the product $\sigma(a_1)\sigma(a_2)$. That is we assign high scores to interaction effects whose corresponding amino acids exhibit large PIPs in our default analysis. We then compute a Spearman correlation coefficient between the PIP ranks for interaction terms in our epistasis analysis with the scores $\sigma(a_1)\sigma(a_2)$. We find a moderate positive correlation given by 0.45. This substantiates our claim that top-scoring interaction effects in our epistasis

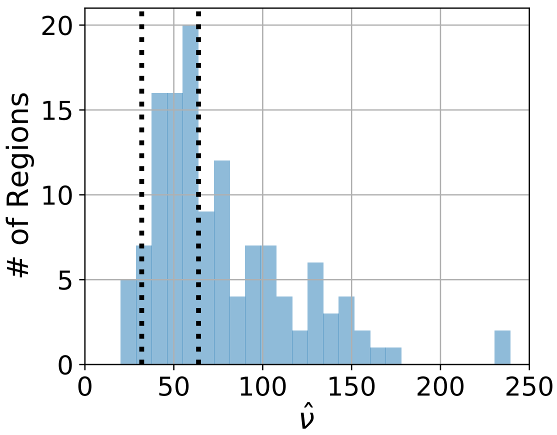


**Fig. S12.** The distribution of estimated effective population sizes $\hat{\nu}$ for the 128 regions in our default analysis. The smallest and largest values of $\hat{\nu}$ are 20.0 and 239.4, respectively. The vertical dotted lines correspond to the median effective population size (63.9; right) and half the median effective population size (32.0; left). The latter quantity is used in our default analysis (see Sec. S6 and Sec. S14.1).

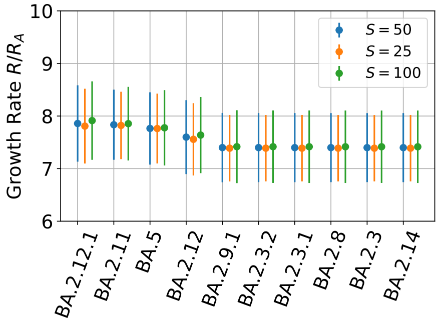


**Fig. S13.** We depict the sensitivity of BVAS estimates of the top 10 lineages w.r.t. the hyperparameter $S$. The results of our default analysis are in blue. Here and elsewhere uncertainty estimates are 95% credible intervals.

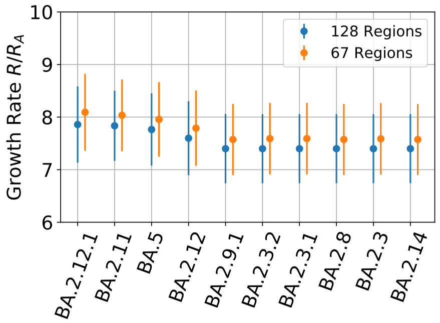


**Fig. S14.** We depict the sensitivity of BVAS estimates of the top 10 lineages w.r.t. the number of regions in the analysis. The results of our default analysis are in blue. Note that the number of regions is effectively controlled by $N_{\rm min}^{\rm tot}$, see Sec. S14.1.

analysis are enriched for interactions between mutations that correspond to top-scoring (linear) effects in our default analysis. To determine the statistical significance of this correlation we conduct a permutation test with 9999 permutations, resulting in a p-value less than 0.0001, i.e. none of the random permutations exceed the observed value of the test statistic.[16]

[16] If we instead use an additive score $\sigma(a_1) + \sigma(a_2)$ we find a correlation of 0.19 and a p-value of $0.0058 \approx 0.006$.

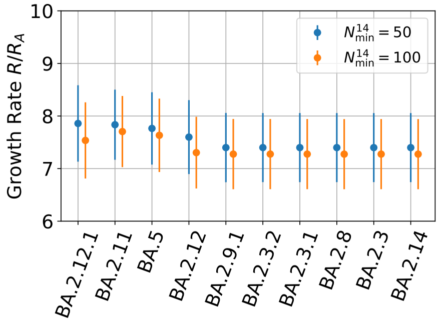


**Fig. S15.** We depict the sensitivity of BVAS estimates of the top 10 lineages w.r.t. the hyperparameter $N_{\rm min}^{14}$ (see Sec. S14.1). The results of our default analysis are in blue.

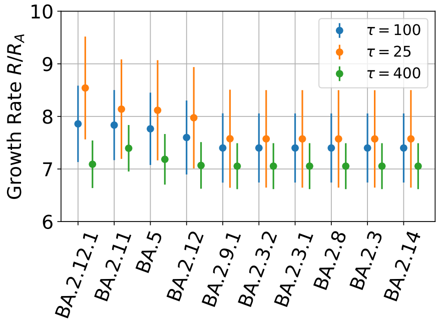


**Fig. S16.** We depict the sensitivity of BVAS estimates of the top 10 lineages w.r.t. the hyperparameter $\tau$, which controls the prior variance of selection coefficients $\beta$. The results of our default analysis are in blue.

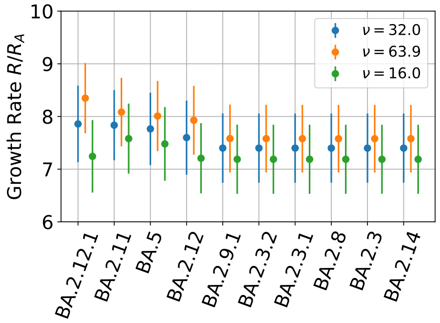


**Fig. S17.** We depict the sensitivity of BVAS estimates of the top 10 lineages w.r.t. three values of effective population size estimate $\hat{\nu}$. The results of our default analysis are in blue. See Sec. S14.1 for details on how we choose $\hat{\nu}$.

### S15. Bayesian variable selection

We briefly review the literature on Bayesian variable selection (BVS). Some of the earliest approaches to BVS were introduced by George and McCulloch (1993, 1997). Chipman et al. (2001) provide an in-depth discussion of BVS for linear regression and CART models. Zanella and Roberts (2019) introduce Tempered Gibbs Sampling (TGS) and apply it to BVS for linear regresion. Dellaportas et al. (2002) and O'Hara et al. (2009) review various methods for BVS.

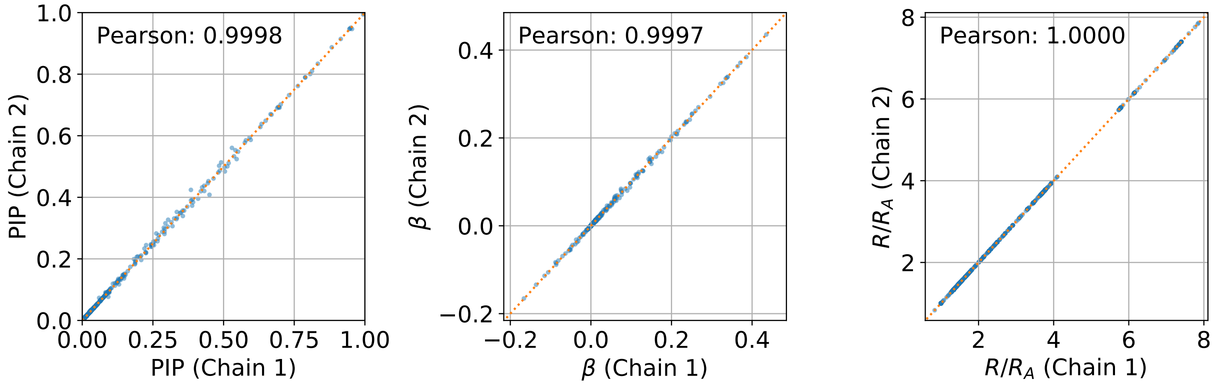


**Fig. S18.** We depict the concordance of BVAS estimates of allele-level Posterior Inclusion Probabilities (left), selection coefficients $\beta$ (middle), and growth rates $R/R_A$ (right) for two analyses run with independent MCMC chains. The near perfect concordance between the two chains suggests that MCMC error is minimal.

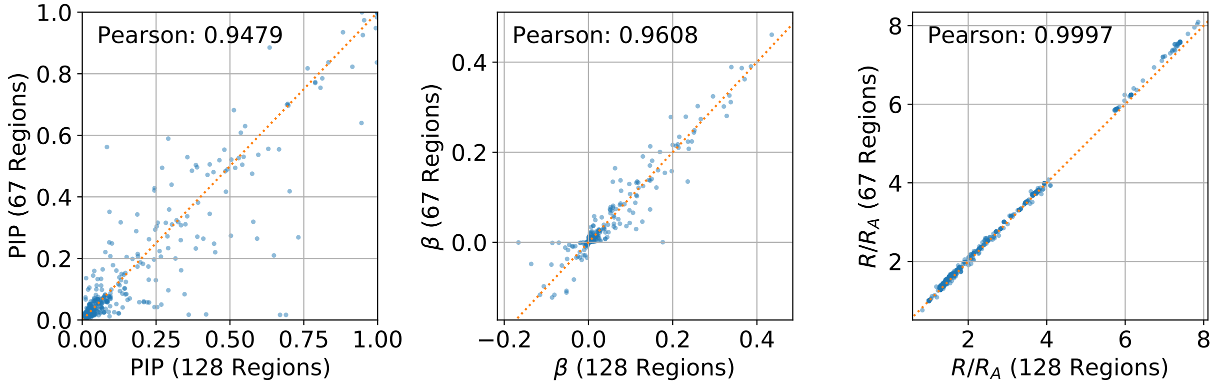


**Fig. S19.** We depict the sensitivity of BVAS estimates of allele-level Posterior Inclusion Probabilities (left), selection coefficients $\beta$ (middle), and growth rates $R/R_A$ (right) for an analysis with the default number of regions ($N_R = 128$) and $N_R = 67$. Note that the number of regions is effectively controlled by $N_{\rm min}^{\rm tot}$, see Sec. S14.1.

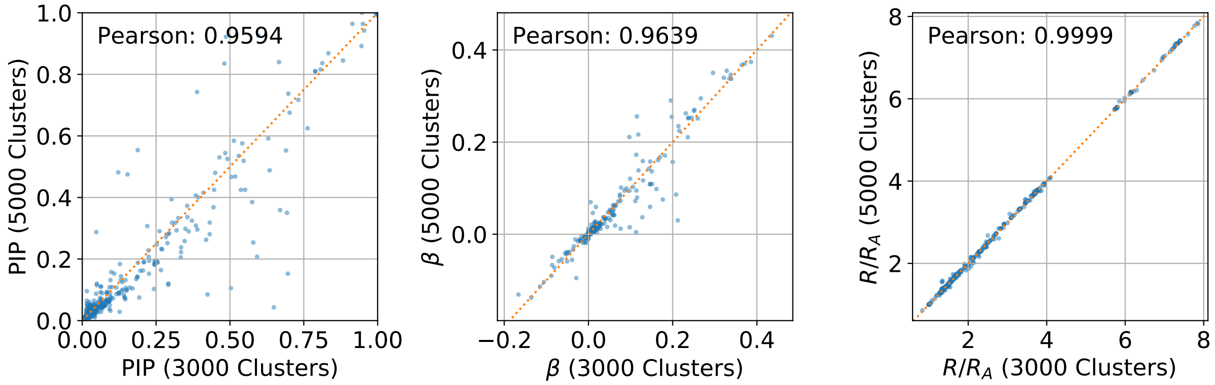


**Fig. S20.** We depict the sensitivity of BVAS estimates of allele-level Posterior Inclusion Probabilities (left), selection coefficients $\beta$ (middle), and growth rates $R/R_A$ (right) for an analysis with the default number of SARS-CoV-2 fine lineage clusters ($L = 3000$) versus $L = 5000$ (see Sec. S14.1). The latter analysis includes $A = 3790$ alleles; here we subset to the $A = 2975$ alleles that are common to both analyses. The top scoring hit that appears in the $A = 3790$ analysis but does not appear in the $A = 2975$ analysis is ORF1a:K1869R with a PIP of 0.068 and $\beta = 0.013$.

| | Region | Raw Counts | Filt. Counts | $\nu$ |
|---|---|---|---|---|
| 1 | United Kingdom / England | 1721377 | 1721335 | 239.4 |
| 2 | USA / California | 477288 | 477275 | 140.9 |
| 3 | United Kingdom / Scotland | 246564 | 241065 | 99.6 |
| 4 | United Kingdom / Wales | 182303 | 176578 | 92.8 |
| 5 | USA / Texas | 179822 | 179814 | 150.9 |
| 6 | Denmark / Hovedstaden | 175112 | 174240 | 92.0 |
| 7 | USA / Colorado | 172317 | 172306 | 131.9 |
| 8 | USA / New York | 140232 | 136337 | 130.4 |
| 9 | USA / Massachusetts | 133818 | 132686 | 128.8 |
| 10 | USA / Florida | 132309 | 132287 | 80.1 |
| 11 | Germany / North Rhine Westphalia | 116236 | 114364 | 147.4 |
| 12 | Canada / British Columbia | 110144 | 109501 | 74.6 |
| 13 | USA / Minnesota | 106425 | 104943 | 136.9 |
| 14 | Germany / Baden Wurttemberg | 88575 | 88158 | 146.7 |
| 15 | USA / Washington | 87015 | 87008 | 79.5 |
| 16 | Denmark / Midtjylland | 82234 | 82167 | 60.0 |
| 17 | USA / Utah | 80841 | 76120 | 154.7 |
| 18 | Denmark / Syddanmark | 77190 | 76982 | 80.8 |
| 19 | USA / Arizona | 73705 | 73701 | 94.4 |
| 20 | USA / Michigan | 70322 | 66930 | 143.7 |
| 21 | USA / North Carolina | 65548 | 64968 | 134.0 |
| 22 | USA / Illinois | 64563 | 64517 | 110.2 |
| 23 | Canada / Ontario | 64419 | 36277 | 25.6 |
| 24 | Turkey | 60669 | 56514 | 50.0 |
| 25 | Germany / Bavaria | 56795 | 56088 | 79.1 |
| 26 | Denmark / Sjaelland | 56097 | 55845 | 75.7 |
| 27 | Brazil / Sao Paulo | 55166 | 53928 | 123.8 |
| 28 | USA / New Jersey | 54349 | 53731 | 101.4 |
| 29 | USA / Wisconsin | 50524 | 49692 | 108.6 |
| 30 | Japan / Tokyo | 49543 | 29498 | 59.9 |
| 31 | USA / Georgia | 48639 | 48016 | 128.2 |
| 32 | USA / Maryland | 48447 | 46665 | 99.1 |
| 33 | USA / Pennsylvania | 47876 | 47184 | 176.1 |
| 34 | Israel | 45954 | 41097 | 72.3 |
| 35 | USA / Tennessee | 45175 | 44716 | 108.0 |
| 36 | USA / Virginia | 43617 | 43608 | 74.6 |
| 37 | France / Auvergne Rhone Alpes | 42853 | 42110 | 86.3 |
| 38 | USA / Ohio | 41591 | 40670 | 167.9 |
| 39 | Germany / Saxony | 40740 | 40666 | 141.3 |
| 40 | Denmark / Nordjylland | 38682 | 38460 | 61.6 |
| 41 | France / Provence Alpes Cote D Azur | 38206 | 35783 | 42.3 |
| 42 | Canada / Alberta | 37606 | 35828 | 42.3 |
| 43 | France / Ile De France | 37591 | 36231 | 92.9 |
| 44 | Canada / Quebec | 36855 | 36850 | 42.4 |
| 45 | Sweden / Stockholm | 36326 | 35446 | 93.8 |
| 46 | USA / Oregon | 36033 | 34458 | 94.8 |
| 47 | USA / Indiana | 34013 | 33602 | 157.7 |
| 48 | USA / Connecticut | 33187 | 32481 | 64.0 |
| 49 | Slovenia | 30151 | 29602 | 65.0 |
| 50 | USA / Nevada | 30092 | 29182 | 99.1 |
| 51 | USA / Wyoming | 29424 | 28919 | 74.6 |
| 52 | Portugal | 29313 | 27052 | 54.9 |
| 53 | South Korea | 28916 | 28655 | 82.1 |
| 54 | Spain / Catalunya | 28673 | 27745 | 55.5 |
| 55 | Italy / Campania | 27781 | 26872 | 46.9 |
| 56 | Australia / Victoria | 27081 | 16390 | 106.8 |
| 57 | Sweden / Skane | 26872 | 26749 | 40.1 |
| 58 | USA / Louisiana | 24383 | 22172 | 48.2 |
| 59 | Australia / New South Wales | 24145 | 21961 | 25.6 |
| 60 | USA / Missouri | 22961 | 22164 | 112.2 |
| 61 | USA / New Mexico | 22845 | 21527 | 91.6 |
| 62 | Luxembourg | 22508 | 22088 | 52.4 |
| 63 | Germany / Hamburg | 21747 | 19962 | 59.2 |
| 64 | France / Nouvelle Aquitaine | 21709 | 21438 | 42.5 |

| | Region | Raw Counts | Filt. Counts | $\nu$ |
|---|---|---|---|---|
| 65 | Japan / Osaka | 21204 | 14531 | 30.9 |
| 66 | USA / Idaho | 21180 | 20850 | 70.8 |
| 67 | Spain / Castilla Y Leon | 20570 | 20065 | 38.4 |
| 68 | USA / Kentucky | 19985 | 19735 | 103.4 |
| 69 | Canada / Saskatchewan | 19674 | 19225 | 49.2 |
| 70 | Germany | 19636 | 19337 | 230.7 |
| 71 | Germany / Schleswig Holstein | 19601 | 17942 | 69.9 |
| 72 | India / Maharashtra | 19130 | 17612 | 58.1 |
| 73 | Germany / Rhineland Palatinate | 18715 | 18675 | 47.6 |
| 74 | Sweden / Vastra Gotaland | 18465 | 16401 | 51.9 |
| 75 | Netherlands / Zuid Holland | 18242 | 17477 | 68.4 |
| 76 | Norway / Viken | 17820 | 17296 | 29.2 |
| 77 | USA / Vermont | 17569 | 16237 | 56.0 |
| 78 | USA / South Carolina | 17378 | 16525 | 125.2 |
| 79 | USA / West Virginia | 17334 | 17274 | 103.9 |
| 80 | USA / Nebraska | 17321 | 16559 | 79.7 |
| 81 | USA / Kansas | 17091 | 16121 | 87.3 |
| 82 | Ireland / Dublin | 16931 | 16109 | 64.4 |
| 83 | Germany / Berlin | 16677 | 16282 | 45.8 |
| 84 | Switzerland / Zurich | 16588 | 10506 | 58.9 |
| 85 | France / Occitanie | 16073 | 11598 | 63.8 |
| 86 | India / Karnataka | 16020 | 15457 | 56.2 |
| 87 | France / Hauts De France | 15960 | 15715 | 56.7 |
| 88 | Switzerland / Bern | 15890 | 12562 | 63.6 |
| 89 | Japan / Saitama | 15856 | 10997 | 30.2 |
| 90 | Japan / Fukuoka | 15152 | 10930 | 20.0 |
| 91 | USA / Maine | 15066 | 14687 | 42.7 |
| 92 | Netherlands / Noord Brabant | 14958 | 14022 | 58.6 |
| 93 | Finland | 14489 | 14416 | 32.4 |
| 94 | France / Grand Est | 14437 | 14255 | 47.2 |
| 95 | Germany / Hesse | 14421 | 14217 | 52.5 |
| 96 | USA / Rhode Island | 14387 | 14159 | 50.2 |
| 97 | Japan | 14251 | 13252 | 25.5 |
| 98 | USA / Montana | 13840 | 13681 | 73.3 |
| 99 | USA / Alabama | 13643 | 13413 | 80.7 |
| 100 | Singapore | 13434 | 11554 | 38.3 |
| 101 | Netherlands / Noord Holland | 13277 | 12893 | 61.0 |
| 102 | USA / Iowa | 13226 | 12665 | 130.2 |
| 103 | Latvia | 13170 | 12470 | 30.8 |
| 104 | Brazil / Rio De Janeiro | 13150 | 12163 | 88.7 |
| 105 | Finland / Uusimaa | 13022 | 10748 | 40.3 |
| 106 | USA / Arkansas | 12974 | 12491 | 43.5 |
| 107 | Japan / Hyogo | 12967 | 6798 | 37.0 |
| 108 | Netherlands / Gelderland | 12955 | 12351 | 59.4 |
| 109 | Switzerland / Geneva | 12952 | 12517 | 45.9 |
| 110 | Norway / Oslo | 12936 | 12403 | 33.2 |
| 111 | Germany / Lower Saxony | 12692 | 12588 | 50.8 |
| 112 | USA / North Dakota | 12462 | 11210 | 71.2 |
| 113 | Mexico / Mexico | 12227 | 11124 | 59.5 |
| 114 | Italy / Lazio | 11918 | 11164 | 40.8 |
| 115 | Switzerland / Basel Landschaft | 11881 | 6588 | 22.3 |
| 116 | Japan / Kanagawa | 11836 | 10546 | 43.1 |
| 117 | Germany / Saxony Anhalt | 11706 | 7713 | 47.9 |
| 118 | USA / Delaware | 11550 | 9082 | 63.0 |
| 119 | Spain / Madrid | 11480 | 10867 | 47.6 |
| 120 | Lithuania / Vilniaus | 11347 | 10199 | 78.1 |
| 121 | Belgium | 11150 | 10368 | 59.7 |
| 122 | Netherlands / Limburg | 11068 | 10789 | 54.0 |
| 123 | Netherlands / Utrecht | 10876 | 10517 | 55.8 |
| 124 | India / Telangana | 10844 | 9522 | 56.0 |
| 125 | Australia / Queensland | 10753 | 9707 | 70.2 |
| 126 | USA / Alaska | 10618 | 10170 | 50.0 |
| 127 | USA / Hawaii | 10348 | 9787 | 45.4 |
| 128 | Belgium / Ghent | 10095 | 9700 | 41.9 |

**Table S2.** Summary statistics for the 128 regions used in our main analyses sorted by raw count totals. All regions are used in our default analysis with $N_{\rm min}^{\rm tot} = 10 \times 10^3$ and the top 67 regions are used in our secondary analysis with $N_{\rm min}^{\rm tot} = 20 \times 10^3$. The 2nd and 3rd columns indicate the total number of samples in each region before and after filtering. The final column denotes the estimated effective population size $\hat{\nu}$. The total number of samples in all regions is 7127962 (raw) and 6910018 (filtered). The total number of samples in the top 67 regions is 6246448 (raw) and 6106650 (filtered). Approximately 3% of the samples are removed in filtering due to the $N_{\rm min}^{14}$ filter (see Sec. S14.1). The average $\hat{\nu}$ in all 128 regions is 77.1; in the top 67 regions it is 92.8. Note that some countries like Germany include both province-level spatial units like Bavaria and larger units that are aggregated from several provinces that are more sparsely sampled (see row 70).

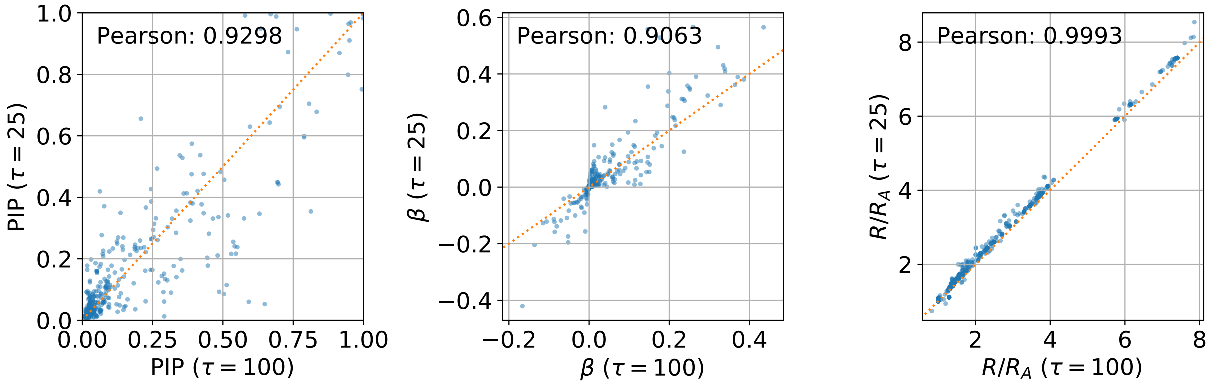


**Fig. S21.** We depict the sensitivity of BVAS estimates of allele-level Posterior Inclusion Probabilities (left), selection coefficients $\beta$ (middle), and growth rates $R/R_A$ (right) for an analysis with the default prior precision ($\tau = 100$) versus $\tau = 25$.

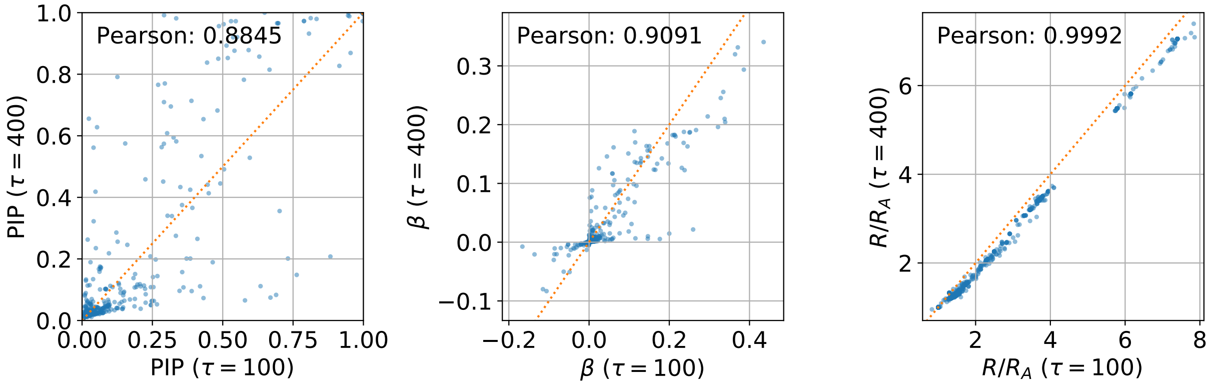


**Fig. S22.** We depict the sensitivity of BVAS estimates of allele-level Posterior Inclusion Probabilities (left), selection coefficients $\beta$ (middle), and growth rates $R/R_A$ (right) for an analysis with the default prior precision ($\tau = 100$) versus $\tau = 400$.

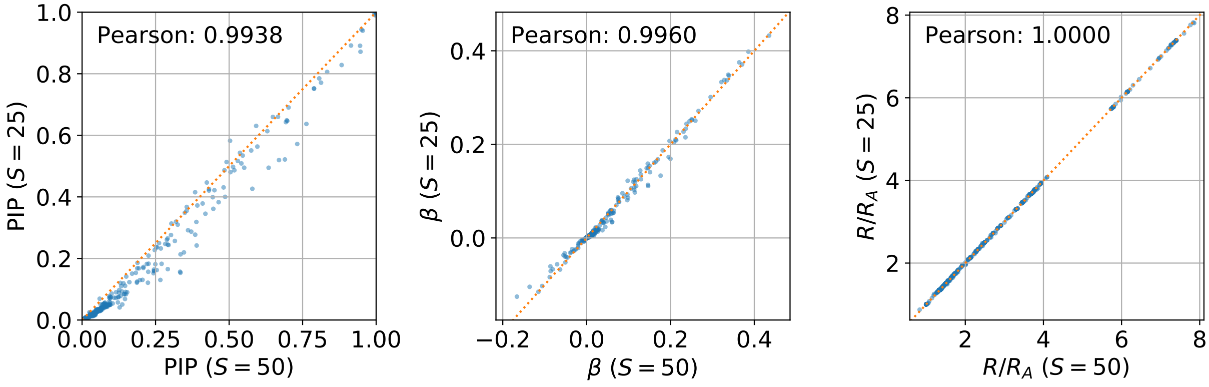


**Fig. S23.** We depict the sensitivity of BVAS estimates of allele-level Posterior Inclusion Probabilities (left), selection coefficients $\beta$ (middle), and growth rates $R/R_A$ (right) for an analysis with the default expected number of non-neutral alleles a priori ($S = 50$) versus $S = 25$. The consequence of halving $S$ is that the total PIP sum decreases from 115.4 to 84.2 and the sum of all PIPs that are larger than 0.1 decreases from 64.2 to 56.4.

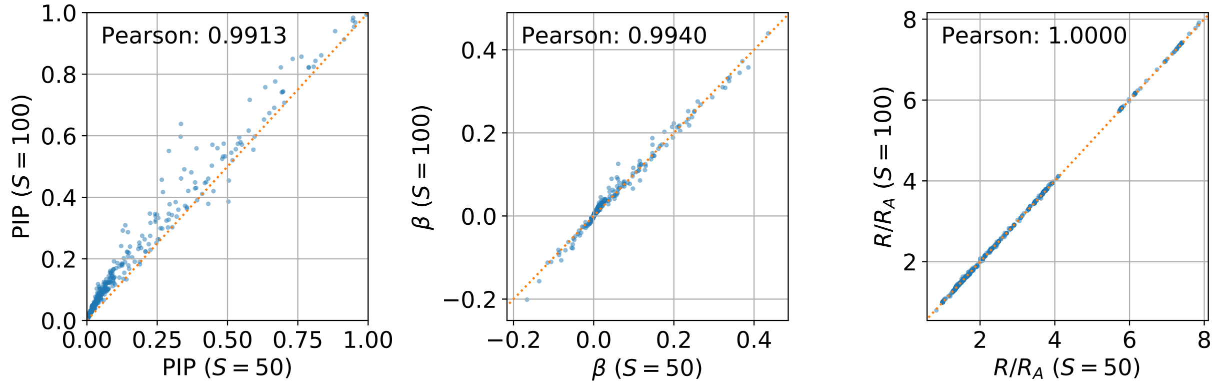


**Fig. S24.** We depict the sensitivity of BVAS estimates of allele-level Posterior Inclusion Probabilities (left), selection coefficients $\beta$ (middle), and growth rates $R/R_A$ (right) for an analysis with the default expected number of non-neutral alleles a priori ($S = 50$) versus $S = 100$. The consequence of doubling $S$ is that the total PIP sum increases from 115.4 to 171.4 and the sum of all PIPs that are larger than 0.1 increases from 64.2 to 78.0.

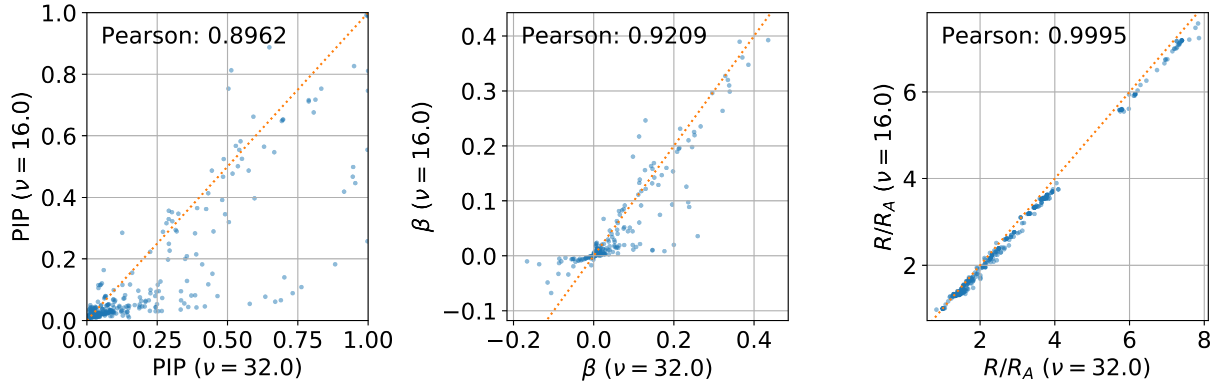


**Fig. S25.** We depict the sensitivity of BVAS estimates of allele-level Posterior Inclusion Probabilities (left), selection coefficients $\beta$ (middle), and growth rates $R/R_A$ (right) for an analysis where the effective population size estimate $\hat{\nu}$ is 16.0 instead of the default 32.0. Notably, there are fewer sizable PIPs in the analysis with smaller $\hat{\nu}$. See Sec. S14.1 for details on how we choose $\hat{\nu}$.

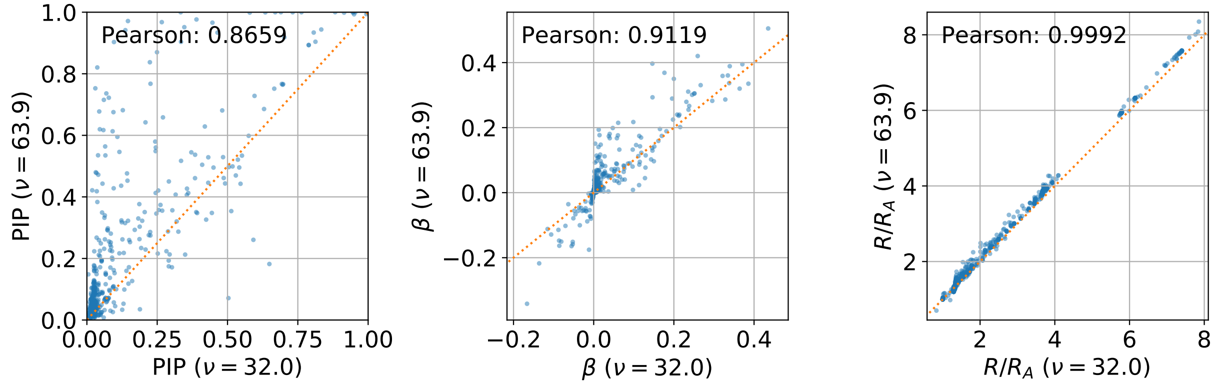


**Fig. S26.** We depict the sensitivity of BVAS estimates of allele-level Posterior Inclusion Probabilities (left), selection coefficients $\beta$ (middle), and growth rates $R/R_A$ (right) for an analysis where the effective population size estimate $\hat{\nu}$ is 63.9 instead of the default 32.0. Notably, there are more sizable PIPs in the analysis with larger $\hat{\nu}$. See Sec. S14.1 for details on how we choose $\hat{\nu}$.

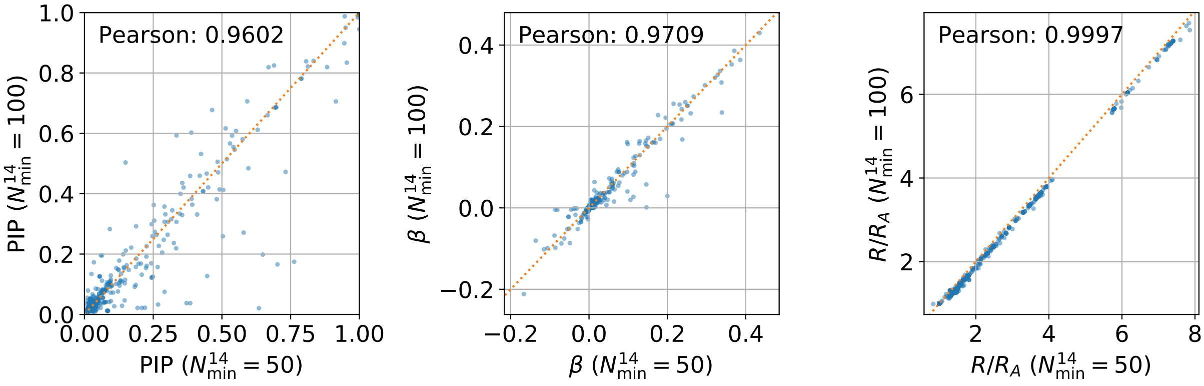


**Fig. S27.** We depict the sensitivity of BVAS estimates of allele-level Posterior Inclusion Probabilities (left), selection coefficients $\beta$ (middle), and growth rates $R/R_A$ (right) for an analysis where the hyperparameter $N^{14}_{\rm min}$ is changed from the default of $N^{14}_{\rm min} = 50$ to $N^{14}_{\rm min} = 100$.

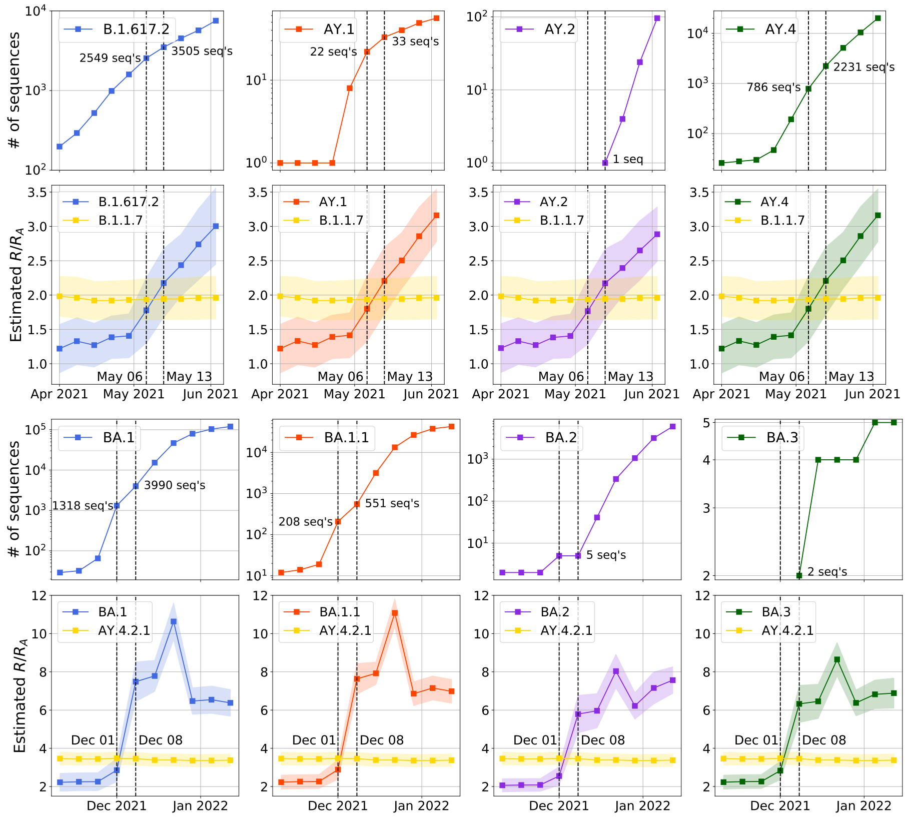


**Fig. S28.** We investigate the ability of BVAS to estimate the fitness of lineages as they emerge. We consider several Delta (top two rows) and Omicron (bottom two rows) sublineages as they emerged in Spring 2021 and the end of 2021, respectively. In both cases we compare to the fittest lineage that was predominant (in England and elsewhere) before the rise of Delta and Omicron. We also depict time series of the number of SARS-CoV-2 sequences in our dataset for each sublineage.

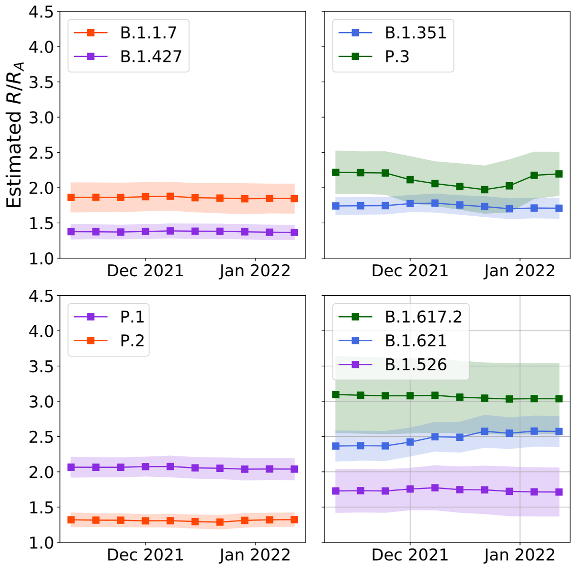


**Fig. S29.** In this companion figure to Figure S28 we depict the stability of growth rate estimates for 9 'well-established' lineages as large numbers of SARS-CoV-2 sequences were collected during the Omicron wave. In particular the total number of sequences included in our analysis increased by about 25%, from 3,901,038 sequences on November 10th to 4,889,119 sequences on January 12th.

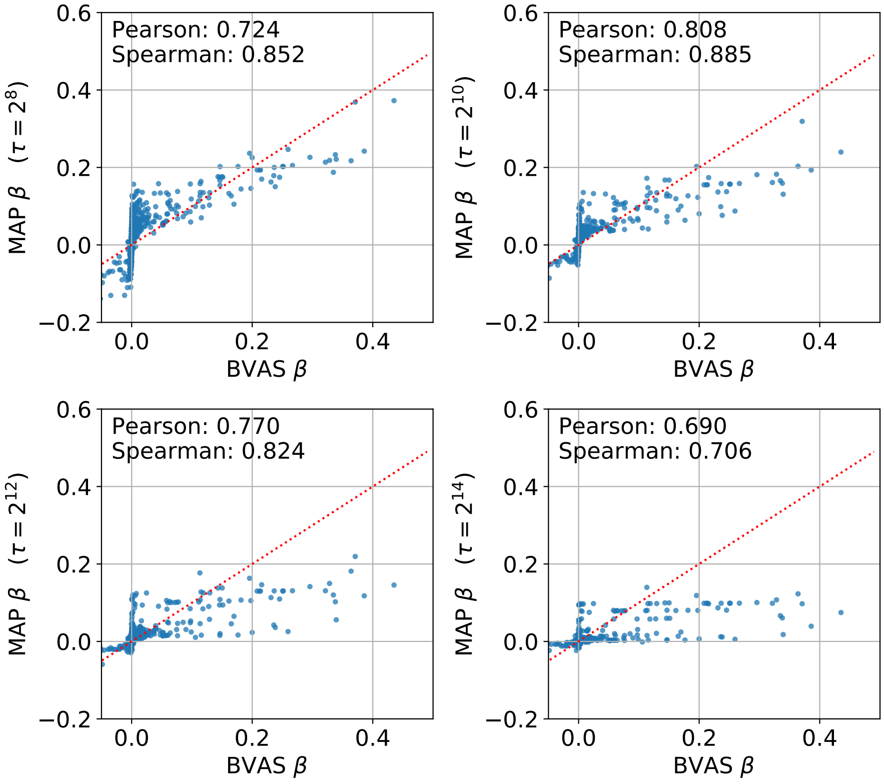


**Fig. S30.** We compare BVAS to MAP estimates for the selection coefficient $\beta$ for 4 values of the MAP regularization parameter $\tau$. As $\tau$ ranges across $\{2^8, 2^{10}, 2^{12}, 2^{14}\}$ the effective regularization parameter $\gamma_{\rm reg} = \tau/\nu$ is approximately given by $\{4, 16, 64, 256\}$, since $\hat{\nu} \approx 63.9$.

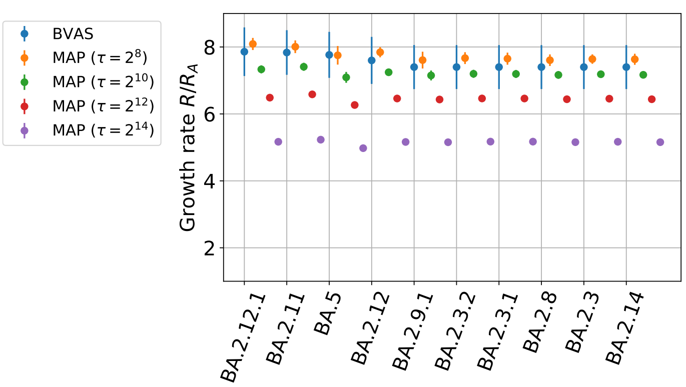


**Fig. S31.** We compare BVAS to MAP estimates for the growth rates of the top 10 fittest lineages (as ranked by BVAS) for 4 values of the MAP regularization parameter $\tau$. MAP growth rate estimates with $\tau \geqslant 10^{12}$ appear to be over-regularized towards Wuhan A. We also note that MAP uncertainty estimates are implausibly narrow.

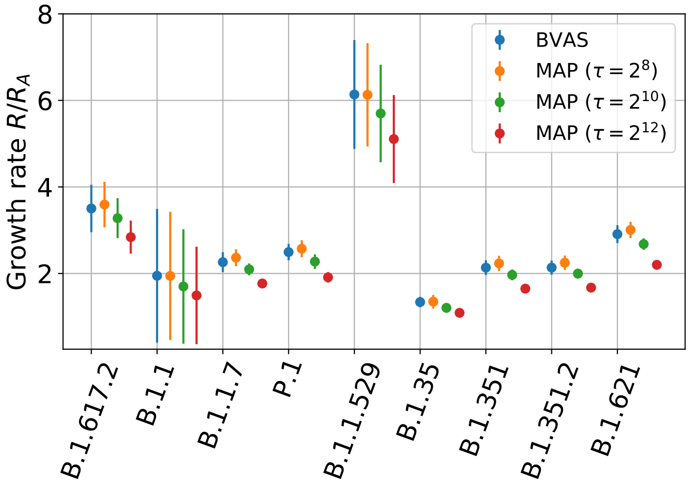


**Fig. S32.** In this companion figure to Figure S31 we compare BVAS to MAP estimates for the growth rates of 9 select lineages for 3 values of the MAP regularization parameter $\tau$. MAP growth rate estimates with $\tau \geqslant 10^{12}$ appear to be over-regularized towards Wuhan A.

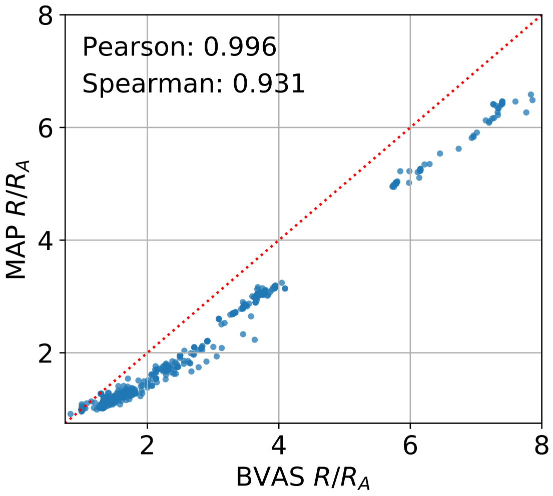


**Fig. S33.** In this companion figure to Figure S31-S32 we compare BVAS and MAP ($\tau = 2^{12}$) estimates of $R/R_A$ across 1662 PANGO lineages. If we regress MAP $R/R_A$ against BVAS $R/R_A$ we find a slope of 0.85, i.e. MAP growth rate estimates are about 15% lower than BVAS growth rate estimates.

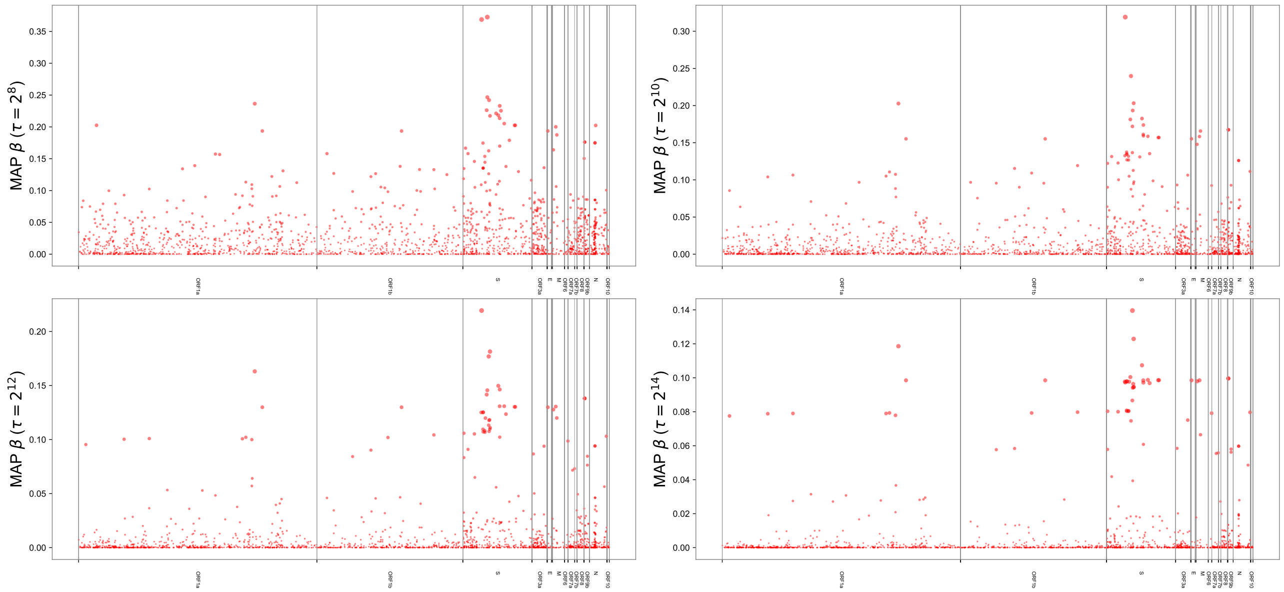


**Fig. S34.** We depict genome-wide Manhattan plots obtained with MAP for 4 distinct values of the regularization parameter $\tau \in \{2^8, 2^{10}, 2^{12}, 2^{14}\}$. These choices of regularization parameters lead to $\{272, 102, 68, 58\}$ alleles with effect sizes greater than 0.05, respectively, and $\{595, 275, 116, 73\}$ alleles with effect sizes greater than 0.025, respectively. MAP results obtained with $\tau \leqslant 2^{10}$ arguably exhibit implausibly high numbers of non-neutral alleles.

| | BVAS Rank | BVAS PIP | BVAS β | MAP Rank | MAP β |
|---|---|---|---|---|---|
| **S:R346K** | 1 | 1.000 | 0.371 ± 0.048 | 1 | 0.219 ± 0.020 |
| **S:L452R** | 2 | 1.000 | 0.435 ± 0.103 | 7 | 0.146 ± 0.027 |
| **S:E484K** | 3 | 1.000 | 0.386 ± 0.097 | 27 | 0.118 ± 0.025 |
| **S:N501Y** | 4 | 1.000 | 0.364 ± 0.118 | 2 | 0.181 ± 0.026 |
| **M:I82T** | 5 | 1.000 | 0.335 ± 0.068 | 24 | 0.120 ± 0.028 |
| **S:D614G** | 6 | 1.000 | 0.339 ± 0.119 | 66 | 0.056 ± 0.029 |
| **S:A222V** | 7 | 1.000 | 0.170 ± 0.058 | 61 | 0.065 ± 0.024 |
| **S:P681H** | 8 | 1.000 | 0.216 ± 0.103 | 6 | 0.146 ± 0.025 |
| **S:S477N** | 9 | 1.000 | 0.231 ± 0.146 | 28 | 0.113 ± 0.026 |
| **S:P681R** | 10 | 1.000 | 0.338 ± 0.146 | 40 | 0.102 ± 0.029 |
| **ORF1b:K1383R** | 11 | 0.999 | -0.106 ± 0.045 | 80 | -0.042 ± 0.022 |
| **S:N440K** | 12 | 0.999 | 0.296 ± 0.126 | 8 | 0.142 ± 0.029 |
| **ORF1b:H1087Y** | 13 | 0.998 | 0.165 ± 0.057 | 77 | 0.046 ± 0.025 |
| **ORF1a:A2529V** | 14 | 0.998 | 0.097 ± 0.039 | 71 | 0.048 ± 0.021 |
| **S:H655Y** | 15 | 0.994 | 0.328 ± 0.154 | 5 | 0.150 ± 0.029 |
| **ORF1a:A2554V** | 16 | 0.954 | -0.115 ± 0.060 | 99 | -0.031 ± 0.023 |
| **S:N679K** | 17 | 0.950 | 0.321 ± 0.233 | 12 | 0.131 ± 0.030 |
| **ORF1a:T3255I** | 18 | 0.947 | 0.196 ± 0.145 | 4 | 0.163 ± 0.026 |
| **S:T859N** | 19 | 0.945 | 0.236 ± 0.182 | 79 | 0.042 ± 0.029 |
| **N:D3L** | 20 | 0.915 | 0.238 ± 0.166 | 87 | 0.036 ± 0.029 |
| **S:L452Q** | 21 | 0.883 | 0.259 ± 0.243 | 124 | 0.025 ± 0.030 |
| **S:N764K** | 22 | 0.833 | 0.267 ± 0.302 | 11 | 0.131 ± 0.030 |
| **M:A63T** | 23 | 0.813 | 0.236 ± 0.263 | 13 | 0.131 ± 0.030 |
| **S:S371F** | 24 | 0.806 | 0.245 ± 0.289 | 31 | 0.110 ± 0.030 |
| **S:N969K** | 25 | 0.789 | 0.250 ± 0.315 | 14 | 0.130 ± 0.030 |

| | BVAS Rank | BVAS PIP | BVAS β | MAP Rank | MAP β |
|---|---|---|---|---|---|
| **S:R346K** | 1 | 1.000 | 0.371 ± 0.048 | 1 | 0.219 ± 0.020 |
| **S:N501Y** | 4 | 1.000 | 0.364 ± 0.118 | 2 | 0.181 ± 0.026 |
| **S:T478K** | 36 | 0.649 | 0.113 ± 0.194 | 3 | 0.177 ± 0.027 |
| **ORF1a:T3255I** | 18 | 0.947 | 0.196 ± 0.145 | 4 | 0.163 ± 0.026 |
| **S:H655Y** | 15 | 0.994 | 0.328 ± 0.154 | 5 | 0.150 ± 0.029 |
| **S:P681H** | 8 | 1.000 | 0.216 ± 0.103 | 6 | 0.146 ± 0.025 |
| **S:L452R** | 2 | 1.000 | 0.435 ± 0.103 | 7 | 0.146 ± 0.027 |
| **S:N440K** | 12 | 0.999 | 0.296 ± 0.126 | 8 | 0.142 ± 0.029 |
| **N:P13L** | 44 | 0.546 | 0.150 ± 0.284 | 9 | 0.138 ± 0.029 |
| **ORF9b:P10S** | 47 | 0.530 | 0.143 ± 0.288 | 10 | 0.138 ± 0.029 |
| **S:N764K** | 22 | 0.833 | 0.267 ± 0.302 | 11 | 0.131 ± 0.030 |
| **S:N679K** | 17 | 0.950 | 0.321 ± 0.233 | 12 | 0.131 ± 0.030 |
| **M:A63T** | 23 | 0.813 | 0.236 ± 0.263 | 13 | 0.131 ± 0.030 |
| **S:N969K** | 25 | 0.789 | 0.250 ± 0.315 | 14 | 0.130 ± 0.030 |
| **S:Q954H** | 26 | 0.789 | 0.252 ± 0.315 | 15 | 0.130 ± 0.030 |
| **ORF1b:I1566V** | 30 | 0.698 | 0.214 ± 0.321 | 16 | 0.130 ± 0.030 |
| **ORF1a:P3395H** | 31 | 0.696 | 0.212 ± 0.321 | 17 | 0.130 ± 0.030 |
| **E:T9I** | 32 | 0.693 | 0.208 ± 0.287 | 18 | 0.130 ± 0.030 |
| **M:Q19E** | 53 | 0.486 | 0.115 ± 0.260 | 19 | 0.128 ± 0.030 |
| **S:S373P** | 54 | 0.484 | 0.119 ± 0.262 | 20 | 0.125 ± 0.030 |
| **S:S375F** | 48 | 0.518 | 0.130 ± 0.266 | 21 | 0.125 ± 0.030 |
| **S:G339D** | 136 | 0.126 | 0.025 ± 0.141 | 22 | 0.125 ± 0.030 |
| **S:D796Y** | 205 | 0.053 | 0.008 ± 0.075 | 23 | 0.124 ± 0.030 |
| **M:I82T** | 5 | 1.000 | 0.335 ± 0.068 | 24 | 0.120 ± 0.028 |
| **S:K417N** | 367 | 0.024 | 0.002 ± 0.030 | 25 | 0.120 ± 0.028 |

**Fig. S35. (Left)** We compare the top 25 BVAS hits to MAP ($\tau = 2^{12}$) hits, where BVAS hits are ranked by PIP. **(Right)** We compare the top 25 MAP ($\tau = 2^{12}$) hits to BVAS hits, where MAP hits are ranked by magnitude of effect size. We note that 12 hits are common to both top 25 rankings.

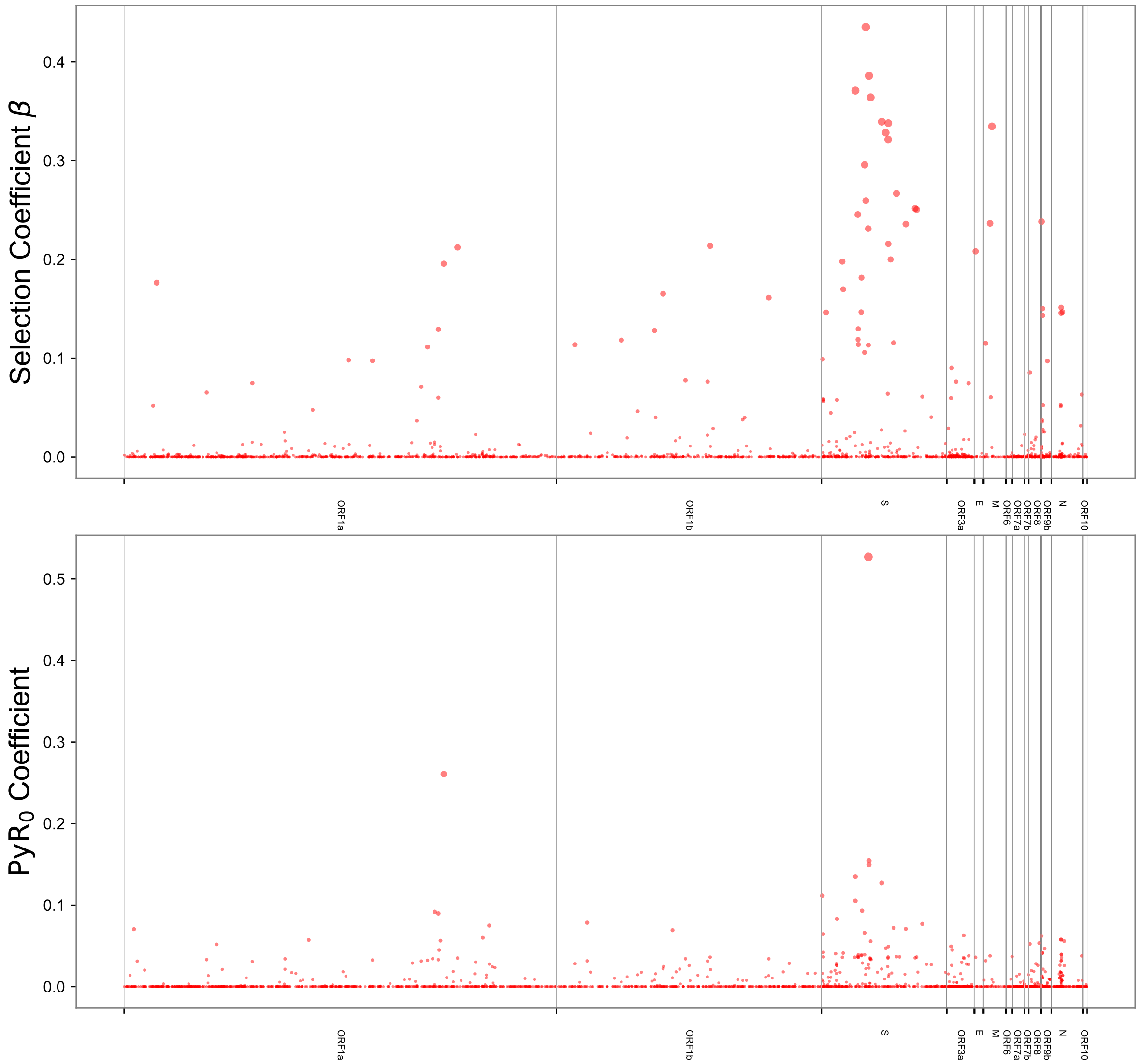


**Fig. S36.** We plot BVAS (top) and $PyR_0$ (bottom) genome-wide Manhattan plots side-by-side. Notably the effects inferred by BVAS are much sparser and the largest effects are larger in magnitude. $PyR_0$ results use a regularization scale $\sigma_3 = 2 \times 10^{-4}$.

| | Alpha PIP | Alpha | Beta PIP | Beta |
|---|---|---|---|---|
| **ORF1a:T265I** | 0.9204 | -0.279 ± 0.218 | 0.0527 | -0.003 ± 0.024 |
| **S:N501Y** | 0.7546 | -0.196 ± 0.263 | 1.0000 | 0.434 ± 0.135 |
| **ORF1a:A2554V** | 0.5340 | -0.097 ± 0.186 | 0.4282 | -0.052 ± 0.120 |
| **ORF3a:T223I** | 0.4853 | 0.118 ± 0.263 | 0.0515 | 0.005 ± 0.048 |
| **ORF1a:A3209V** | 0.4342 | -0.094 ± 0.230 | 0.0068 | 0.000 ± 0.010 |

**Table S3.** In this companion table to Table 3 in the main text we report PIPs and estimated coefficients $\alpha$ for vaccination-dependent effects for an analysis in which the vaccination status is 'vaccinated'. We report effects that are ranked in the top 75 by PIP. Estimated $\beta$ coefficients and PIPs for the corresponding vaccination-independent effects are reported in the two rightmost columns.

| | BVAS PIP | BVAS β | BVAS Rank | PyR0 Coef. | PyR0 Rank |
|---|---|---|---|---|---|
| S:R346K | 1.000 | 0.371 ± 0.048 | 1 | 0.105 ± 0.00007 | 5 |
| S:L452R | 1.000 | 0.435 ± 0.103 | 2 | 0.022 ± 0.00003 | 44 |
| S:E484K | 1.000 | 0.386 ± 0.097 | 3 | 0.155 ± 0.00042 | 50 |
| S:N501Y | 1.000 | 0.364 ± 0.118 | 4 | 0.056 ± 0.00004 | 2 |
| M:I82T | 1.000 | 0.335 ± 0.068 | 5 | 0.009 ± 0.00003 | 53 |
| S:D614G | 1.000 | 0.339 ± 0.119 | 6 | 0.127 ± 0.00107 | 84 |
| S:A222V | 1.000 | 0.170 ± 0.058 | 7 | 0.041 ± 0.00009 | 48 |
| S:P681H | 1.000 | 0.216 ± 0.103 | 8 | 0.049 ± 0.00004 | 10 |
| S:S477N | 1.000 | 0.231 ± 0.146 | 9 | 0.027 ± 0.00003 | 38 |
| S:P681R | 1.000 | 0.338 ± 0.146 | 10 | 0.012 ± 0.00003 | 49 |
| ORF1b:K1383R | 0.999 | -0.106 ± 0.045 | 11 | -0.037 ± 0.00073 | 139 |
| S:N440K | 0.999 | 0.296 ± 0.126 | 12 | 0.039 ± 0.00003 | 30 |
| ORF1b:H1087Y | 0.998 | 0.165 ± 0.057 | 13 | 0.022 ± 0.00020 | 89 |
| ORF1a:A2529V | 0.998 | 0.097 ± 0.039 | 14 | 0.033 ± 0.00012 | 59 |
| S:H655Y | 0.994 | 0.328 ± 0.154 | 15 | 0.047 ± 0.00003 | 1 |
| ORF1a:A2554V | 0.954 | -0.115 ± 0.060 | 16 | -0.003 ± 0.00011 | 184 |
| S:N679K | 0.950 | 0.321 ± 0.233 | 17 | 0.036 ± 0.00003 | 16 |
| ORF1a:T3255I | 0.947 | 0.196 ± 0.145 | 18 | 0.261 ± 0.00035 | 45 |
| S:T859N | 0.945 | 0.236 ± 0.182 | 19 | 0.071 ± 0.00094 | 117 |
| N:D3L | 0.915 | 0.238 ± 0.166 | 20 | 0.062 ± 0.00025 | 61 |
| S:L452Q | 0.883 | 0.259 ± 0.243 | 21 | 0.001 ± 0.00088 | 380 |
| S:N764K | 0.833 | 0.267 ± 0.302 | 22 | 0.037 ± 0.00003 | 34 |
| M:A63T | 0.813 | 0.236 ± 0.263 | 23 | 0.038 ± 0.00003 | 18 |
| S:S371F | 0.806 | 0.245 ± 0.289 | 24 | 0.037 ± 0.00003 | 20 |
| S:N969K | 0.789 | 0.250 ± 0.315 | 25 | 0.036 ± 0.00003 | 31 |

| | BVAS PIP | BVAS β | BVAS Rank | PyR0 Coef. | PyR0 Rank |
|---|---|---|---|---|---|
| S:H655Y | 0.994 | 0.328 ± 0.154 | 15 | 0.047 ± 0.00003 | 1 |
| S:N501Y | 1.000 | 0.364 ± 0.118 | 4 | 0.056 ± 0.00004 | 2 |
| S:V213G | 0.666 | 0.198 ± 0.316 | 35 | 0.036 ± 0.00002 | 3 |
| S:D405N | 0.542 | 0.147 ± 0.299 | 45 | 0.038 ± 0.00003 | 4 |
| S:R346K | 1.000 | 0.371 ± 0.048 | 1 | 0.105 ± 0.00007 | 5 |
| S:T376A | 0.444 | 0.114 ± 0.287 | 59 | 0.039 ± 0.00003 | 6 |
| N:P13L | 0.546 | 0.150 ± 0.284 | 44 | 0.041 ± 0.00003 | 7 |
| ORF1a:T3090I | 0.431 | 0.111 ± 0.278 | 61 | 0.032 ± 0.00002 | 8 |
| ORF1b:T2163I | 0.575 | 0.161 ± 0.300 | 42 | 0.034 ± 0.00002 | 9 |
| S:P681H | 1.000 | 0.216 ± 0.103 | 8 | 0.049 ± 0.00004 | 10 |
| ORF3a:T223I | 0.153 | 0.018 ± 0.090 | 120 | 0.038 ± 0.00003 | 11 |
| ORF1b:R1315C | 0.302 | 0.078 ± 0.247 | 83 | 0.034 ± 0.00002 | 12 |
| S:R408S | 0.630 | 0.181 ± 0.305 | 38 | 0.039 ± 0.00003 | 13 |
| S:S373P | 0.484 | 0.119 ± 0.262 | 54 | 0.036 ± 0.00003 | 14 |
| N:S413R | 0.291 | 0.063 ± 0.192 | 86 | 0.038 ± 0.00003 | 15 |
| S:N679K | 0.950 | 0.321 ± 0.233 | 17 | 0.036 ± 0.00003 | 16 |
| ORF9b:P10S | 0.530 | 0.143 ± 0.288 | 47 | 0.041 ± 0.00003 | 17 |
| M:A63T | 0.813 | 0.236 ± 0.263 | 23 | 0.038 ± 0.00003 | 18 |
| S:G339D | 0.126 | 0.025 ± 0.141 | 136 | 0.036 ± 0.00003 | 19 |
| S:S371F | 0.806 | 0.245 ± 0.289 | 24 | 0.037 ± 0.00003 | 20 |
| ORF6:D61L | 0.041 | 0.004 ± 0.049 | 231 | 0.037 ± 0.00003 | 21 |
| S:Q493R | 0.041 | 0.006 ± 0.064 | 232 | 0.035 ± 0.00003 | 22 |
| ORF1a:L3201F | 0.316 | 0.060 ± 0.188 | 81 | 0.033 ± 0.00003 | 23 |
| S:D796Y | 0.053 | 0.008 ± 0.075 | 205 | 0.036 ± 0.00003 | 24 |
| ORF1a:P3395H | 0.696 | 0.212 ± 0.321 | 31 | 0.035 ± 0.00003 | 25 |

**Fig. S37. (Left)** We compare the top 25 BVAS hits to $\text{PyR}_0$ hits, where BVAS hits are ranked by PIP. **(Right)** We compare the top 25 $\text{PyR}_0$ hits to BVAS hits, where $\text{PyR}_0$ hits are ranked by z-score. We note that $\text{PyR}_0$ uncertainty estimates are implausibly small, an unfortunate consequence of variational inference. We note that 7 hits are common to both top 25 rankings.

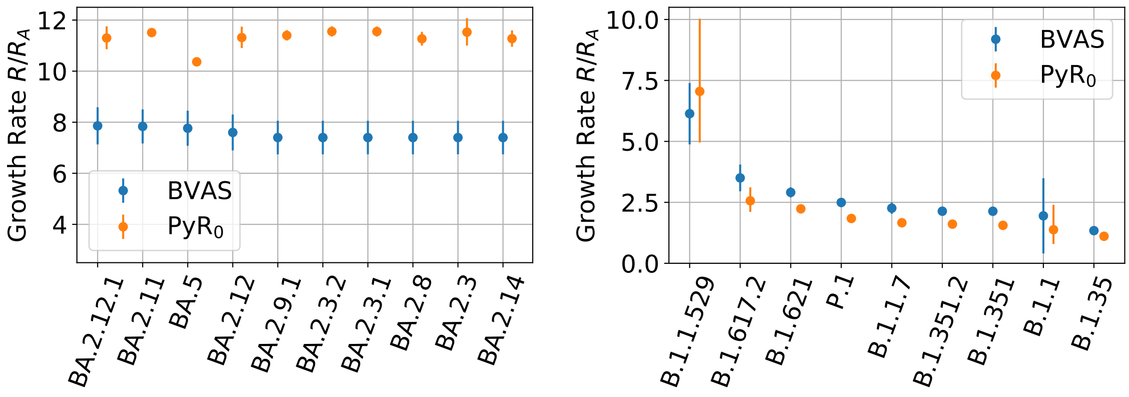


**Fig. S38.** We compare BVAS and $\text{PyR}_0$ estimates of $R/R_A$. Uncertainty bars denote 95% credibile intervals. **(Left)** We compare the top 10 most fit lineages (as ranked by BVAS). **(Right)** We compare a selection of notable lineages (as ranked by BVAS). $\text{PyR}_0$ uncertainty estimates tend to be much narrower, a property that results from overconfident uncertainty estimates of mutation-specific coefficients. Note that both methods infer significant diversity in B.1.1 fitness estimates (see Figure S7).

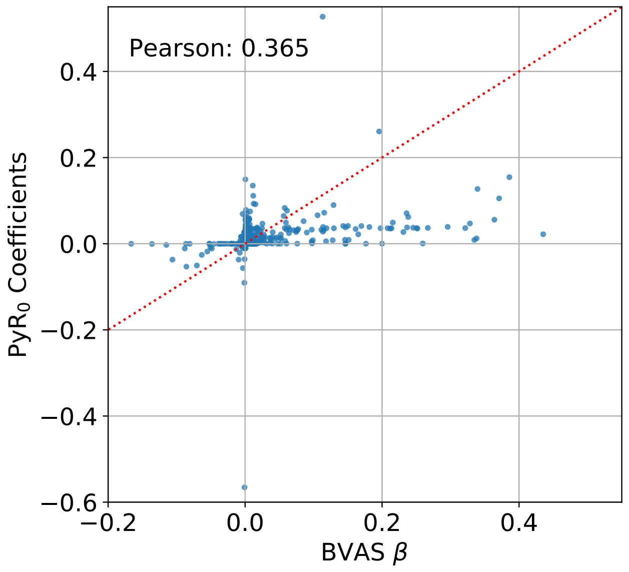


**Fig. S39.** We plot $\beta$ coefficients inferred by BVAS against coefficients inferred by $PyR_0$. Notably $PyR_0$ infers many more non-negligible coefficients. As a consequence the effect sizes of top BVAS hits tend to be considerably larger. $PyR_0$ results use a regularization scale $\sigma_3 = 2 \times 10^{-4}$.

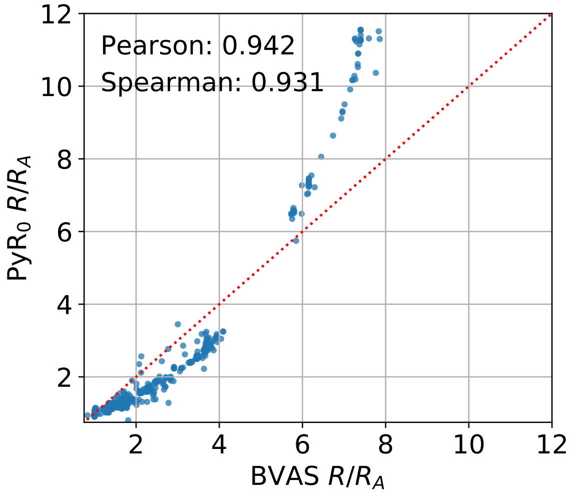


**Fig. S40.** We compare BVAS and $PyR_0$ estimates of $R/R_A$ across 1662 PANGO lineages. $PyR_0$ estimates are markedly higher than BVAS estimates for Omicron lineages and otherwise tend to be lower.

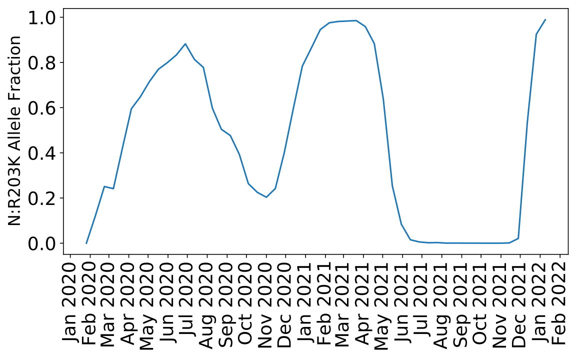


**Fig. S41.** We depict allele fraction time series for N:R203K in England throughout the first two years of the pandemic. The allele fractions for ORF14:G50N and N:G204R are nearly identical. BVAS infers that all three alleles are likely passenger mutations.

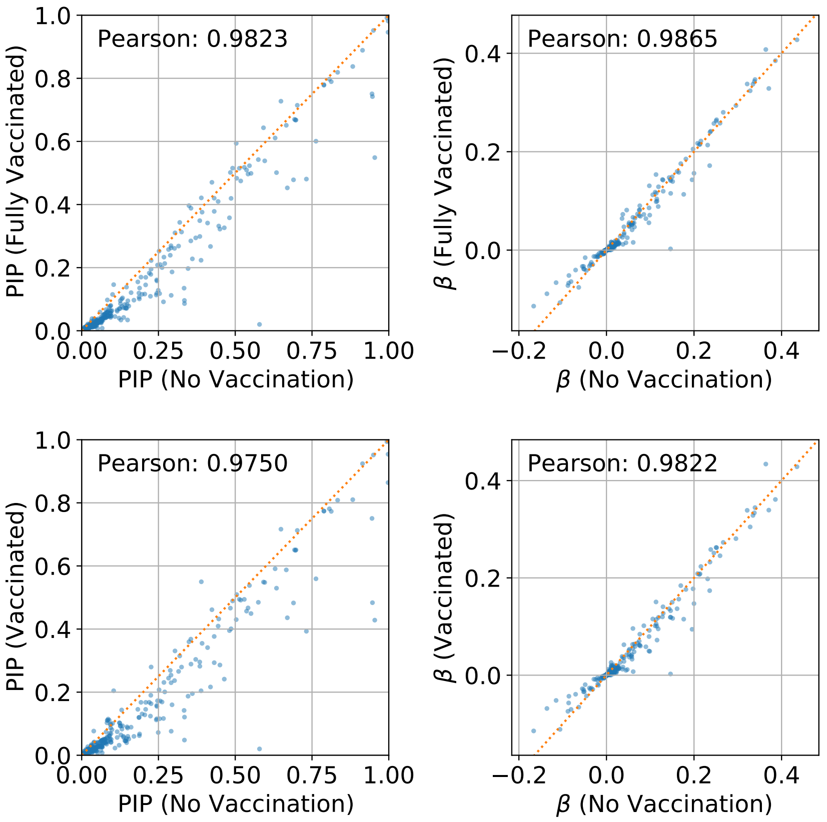


**Fig. S42.** We compare BVAS estimates from vaccination-independent analyses to estimates derived from analyses with vaccination-dependent effects. We depict allele-level Posterior Inclusion Probabilities (left) and selection coefficients $\beta$ (right); i.e. both quantitites correspond to vaccination-independent effects. The vaccination status used in the top (respectively, bottom) panel is 'fully vaccinated' ('vaccinated'). In both panels $N_{\rm min}^{\rm tot} = 10^4$ and $N_R = 127$ regions are included in the analysis. The strong concordance implies that the incorporation of vaccination status into the model has only moderate effects on the estimates of vaccination-independent effects and model fit as a whole.

| | PIP | AA1 | AA2 | AA1 Counts | AA2 Counts | AA1+AA2 Counts | AA1 PANGOs | AA2 PANGOs | AA1+AA2 PANGOs | Quad. Beta12 | Quad. Beta1 | Quad. Beta2 | Lin. Beta1 | Lin. Beta2 |
|---|---|---|---|---|---|---|---|---|---|---|---|---|---|---|
| **1** | 0.811 | N501Y | P681H | 155918 | 61153 | 1343290 | P.1, B.1.351 | B.1.1.519, B.1.243 | B.1.1.7, BA.1 | 0.2016 | 0.3659 | 0.1064 | 0.3640 | 0.2157 |
| **2** | 0.528 | P681H | T1027I | 1394250 | 113678 | 10193 | B.1.1.7, BA.1 | P.1, P.1.14 | P.1.7, XB | 0.1403 | 0.1064 | 0.0906 | 0.2157 | 0.0611 |
| **3** | 0.359 | A67V | E484A | 11580 | 6778 | 184464 | B.1.525, BA.1 | BA.2, B.1.633 | BA.1, BA.1.1 | -0.0846 | -0.0004 | 0.0033 | -0.0023 | 0.0003 |
| **4** | 0.270 | L452R | S12F | 3885747 | 2880 | 5761 | AY.4, AY.103 | B.1.1.63 | AY.39.1, C.36.3 | 0.0597 | 0.4473 | 0.0781 | 0.4352 | 0.0988 |
| **5** | 0.269 | N501Y | A67V | 1317045 | 13881 | 182163 | B.1.1.7, P.1 | BA.1, B.1.525 | BA.1, BA.1.1 | -0.0638 | 0.3659 | -0.0004 | 0.3640 | -0.0023 |
| **6** | 0.263 | N501Y | H49Y | 1498176 | 1290 | 1032 | B.1.1.7, BA.1 | B.1.369, AY.33.1 | P.1.17.1, B.1.604 | 0.0614 | 0.3659 | 0.1283 | 0.3640 | 0.1464 |
| **7** | 0.209 | L452R | T478K | 103192 | 214285 | 3788316 | B.1.429, B.1.427 | BA.1, BA.1.1 | AY.4, AY.103 | 0.0502 | 0.4473 | 0.2015 | 0.4352 | 0.1132 |
| **8** | 0.203 | P681H | N440K | 1214913 | 7127 | 189530 | B.1.1.7, B.1.1.519 | B.1.36.29, B.1.619.1 | BA.1, BA.1.1 | 0.0495 | 0.1064 | 0.2637 | 0.2157 | 0.2958 |

**Table S4.** We report inference results for the top hits in our epistasis analysis, namely those with PIPs above 0.2. These are the same hits that are depicted in the interaction network diagram in the main text. The first column reports the PIP of the pairwise interaction term in the epistasis analysis with both linear and quadratic effects. The next two columns list the corresponding Spike amino acid mutations. The next two columns report the total number of SARS-CoV-2 sequences in our dataset that carry the first (respectively, second) but not the second (respectively, first) amino acid mutation. The next column reports the total number of SARS-CoV-2 sequences in our dataset that carry both amino acid mutations. The next three columns are like the counts in the previous three columns except they report the leading PANGO lineages corresponding to the counts in the previous three columns. Note that a lineage can appear in multiple columns because there is non-negligible genetic diversity in some PANGO lineages (e.g. BA.1). The next column reports the inferred interaction effect in the epistasis analysis with both linear and quadratic effects. The next two columns report the inferred linear effects in the epistasis analysis corresponding to amino acids AA1 and AA2. The last two columns report the inferred linear effects in the default analysis corresponding to amino acids AA1 and AA2.

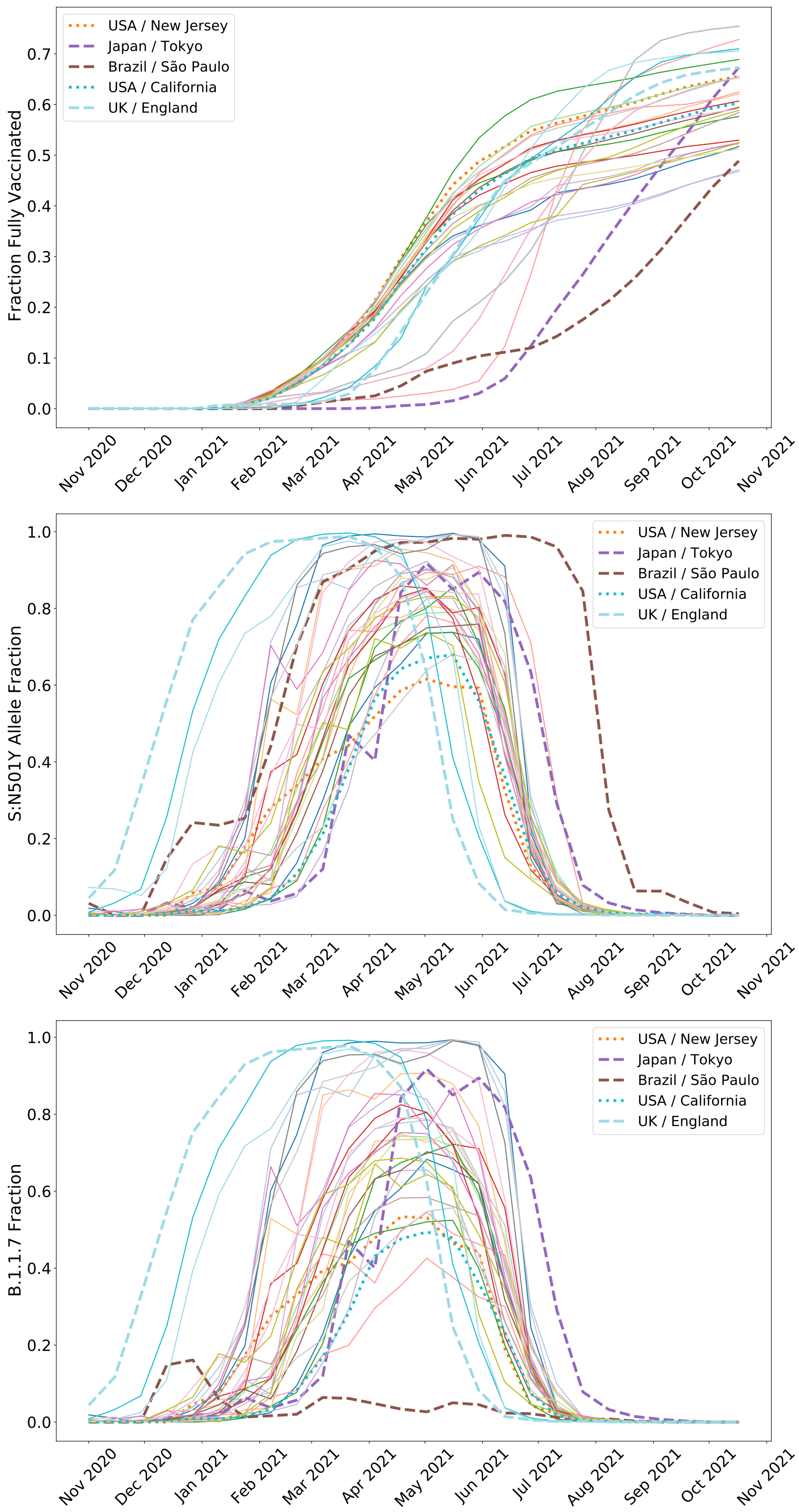


**Fig. S43. (Top)** We depict time series for the fraction of individuals who are fully vaccinated for 34 well-sampled regions (Ritchie et al., 2020). **(Middle)** We depict allele fractions for the N501Y mutation in the S gene for the same 34 regions. **(Bottom)** We depict variant fractions for the B.1.1.7 (Alpha) variant for the same 34 regions. The rise and fall of B.1.1.7 is correlated with that of S:N501Y but not identical; for example, B.1.1.7 never became dominant in Brazil.

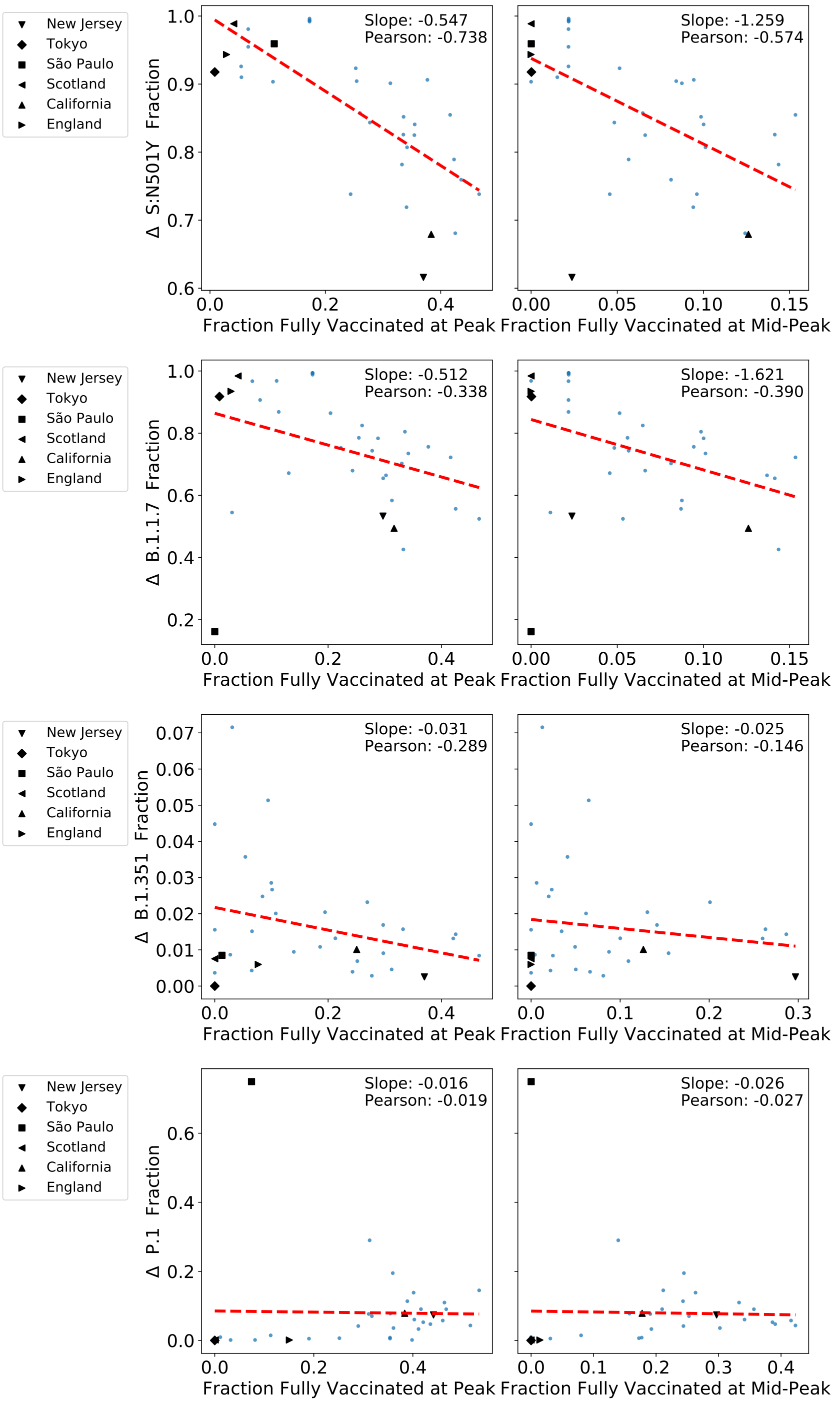


**Fig. S44.** We correlate vaccination status with uptake (i.e. rising prevalance) of the S:N501Y allele as well as uptake of 3 SARS-CoV-2 variants that contain S:N501Y. In each case the vertical axis encodes the difference in allele/variant fraction from November 1$^{st}$ 2020 to the subsequent peak (typically in March to June 2021). The horizontal axis depicts the vaccination status at the peak or at mid-peak, i.e. when the corresponding allele/variant fraction has obtained half the value of the peak. Red dashed lines depict linear fits.

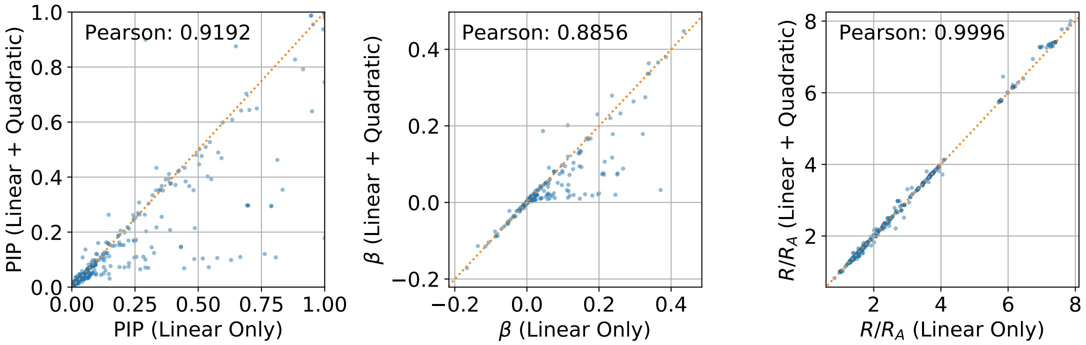


**Fig. S45.** We depict the sensitivity of BVAS estimates of allele-level Posterior Inclusion Probabilities (left), selection coefficients $\beta$ (middle), and growth rates $R/R_A$ (right) for an analysis with linear and quadratic effects to one with linear effects only. The former analysis includes 4407 selection effects; here we subset to the $A = 2975$ linear effects that are common to both analyses. The inferred linear effects are largely the same for both analyses, except for a relatively modest number of outliers. In particular there are 5 linear effects whose PIP is substantially smaller in the epistasis analysis (S:R346K, S:S371F, S:S704L, S:V213G, and S:R408S) and one linear effect (S:T95I) whose PIP is substantially larger in the epistasis analysis (where a substantial difference means a difference larger than 0.5). As expected growth rate estimates exhibit very high concordance.

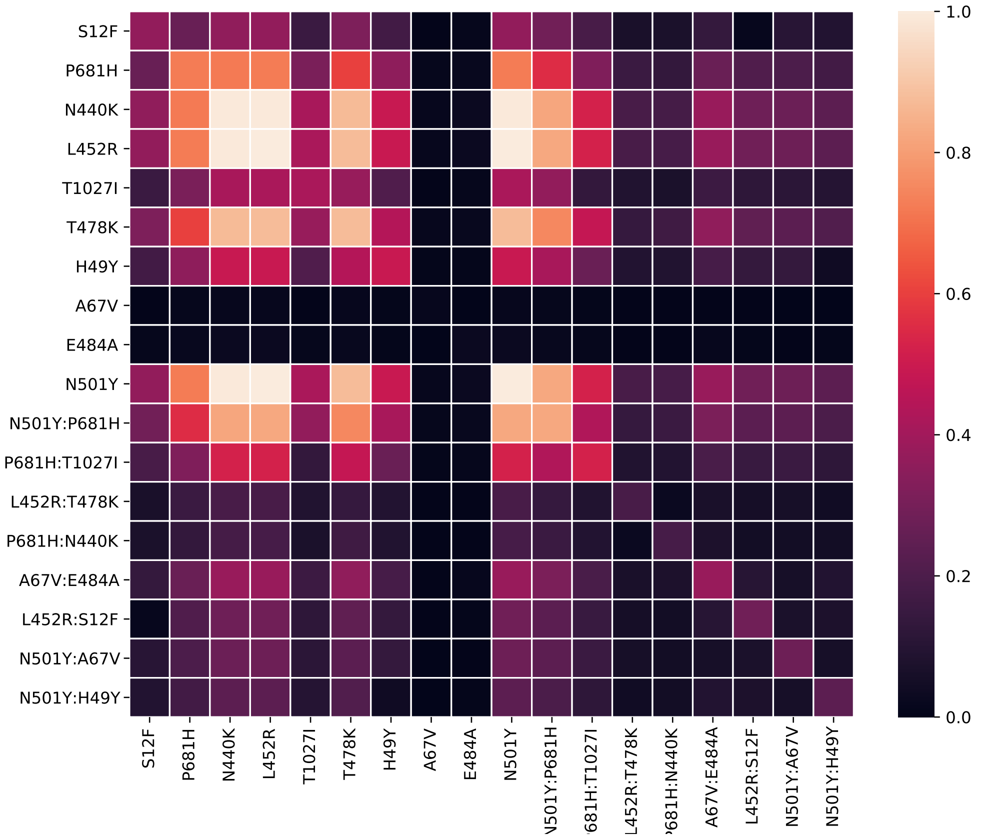


**Fig. S46.** We depict co-occurrence posterior inclusion probabilities for the 8 interaction effects and corresponding 10 linear effects in our epistasis analysis. In other words diagonal entries correspond to standard PIPs, while off-diagonal entries correspond to the posterior probability that the corresponding *pair* of features co-occur in the model posterior. For example the interaction term (N501Y, P681H) co-occurs frequently with (P681H, T1027I) in the posterior.

### References

Q. Bi, Y. Wu, S. Mei, C. Ye, X. Zou, Z. Zhang, X. Liu, L. Wei, S. A. Truelove, T. Zhang, et al. Epidemiology and transmission of covid-19 in 391 cases and 1286 of their close contacts in shenzhen, china: a retrospective cohort study. *The Lancet infectious diseases*, 20(8):911–919, 2020.

E. Bingham, J. P. Chen, M. Jankowiak, F. Obermeyer, N. Pradhan, T. Karaletsos, R. Singh, P. A. Szerlip, P. Horsfall, and N. D. Goodman. Pyro: Deep universal probabilistic programming. *J. Mach. Learn. Res.*, 20:28:1–28:6, 2019. URL `http://jmlr.org/papers/v20/18-403.html`.

P. G. Bissiri, C. C. Holmes, and S. G. Walker. A general framework for updating belief distributions. *Journal of the Royal Statistical Society: Series B (Statistical Methodology)*, 78(5):1103–1130, 2016.

C. M. Carvalho, N. G. Polson, and J. G. Scott. Handling sparsity via the horseshoe. In *Artificial Intelligence and Statistics*, pages 73–80. PMLR, 2009.

E. R. Chan, L. D. Jones, M. Linger, J. D. Kovach, M. M. Torres-Teran, A. Wertz, C. J. Donskey, and P. A. Zimmerman. Covid-19 infection and transmission includes complex sequence diversity. *bioRxiv*, 2022.

B. Charlesworth, M. Morgan, and D. Charlesworth. The effect of deleterious mutations on neutral molecular variation. *Genetics*, 134(4):1289–1303, 1993.

Z. Chen, D. Pei, L. Jiang, Y. Song, J. Wang, H. Wang, D. Zhou, J. Zhai, Z. Du, B. Li, et al. Antigenicity analysis of different regions of the severe acute respiratory syndrome coronavirus nucleocapsid protein. *Clinical chemistry*, 50(6):988–995, 2004.

H. Chipman, E. I. George, R. E. McCulloch, M. Clyde, D. P. Foster, and R. A. Stine. The practical implementation of bayesian model selection. *Lecture Notes-Monograph Series*, pages 65–134, 2001.

J. Cubuk, J. J. Alston, J. J. Incicco, S. Singh, M. D. Stuchell-Brereton, M. D. Ward, M. I. Zimmerman, N. Vithani, D. Griffith, J. A. Wagoner, et al. The sars-cov-2 nucleocapsid protein is dynamic, disordered, and phase separates with rna. *Nature communications*, 12(1):1–17, 2021.

P. Dellaportas, J. J. Forster, and I. Ntzoufras. On bayesian model and variable selection using mcmc. *Statistics and Computing*, 12 (1):27–36, 2002.

S. Elbe and G. Buckland-Merrett. Data, disease and diplomacy: Gisaid's innovative contribution to global health. *Global challenges*, 1(1):33–46, 2017.

A. Endo et al. Estimating the overdispersion in covid-19 transmission using outbreak sizes outside china. *Wellcome open research*, 5, 2020.

E. I. George and R. E. McCulloch. Variable selection via gibbs sampling. *Journal of the American Statistical Association*, 88(423): 881–889, 1993.

E. I. George and R. E. McCulloch. Approaches for bayesian variable selection. *Statistica sinica*, pages 339–373, 1997.

A. J. Greaney, T. N. Starr, and J. D. Bloom. An antibody-escape estimator for mutations to the sars-cov-2 receptor-binding domain. *Virus Evolution*, 2022.

M. Jankowiak. Fast bayesian variable selection in binomial and negative binomial regression. *arXiv preprint arXiv:2106.14981*, 2021.

D. P. Kingma and J. Ba. Adam: A method for stochastic optimization. *arXiv preprint arXiv:1412.6980*, 2014.

M. S. Lau, B. Grenfell, M. Thomas, M. Bryan, K. Nelson, and B. Lopman. Characterizing superspreading events and age-specific infectiousness of sars-cov-2 transmission in georgia, usa. *Proceedings of the National Academy of Sciences*, 117(36):22430–22435, 2020.

B. Lee, M. S. Sohail, E. Finney, S. F. Ahmed, A. A. Quadeer, M. R. McKay, and J. P. Barton. Inferring effects of mutations on sars-cov-2 transmission from genomic surveillance data. *medRxiv*, pages 2021–12, 2022.

J. McBroome, B. Thornlow, A. S. Hinrichs, A. Kramer, N. De Maio, N. Goldman, D. Haussler, R. Corbett-Detig, and Y. Turakhia. A daily-updated database and tools for comprehensive sars-cov-2 mutation-annotated trees. *Molecular biology and evolution*, 38 (12):5819–5824, 2021.

D. Miller, M. A. Martin, N. Harel, O. Tirosh, T. Kustin, M. Meir, N. Sorek, S. Gefen-Halevi, S. Amit, O. Vorontsov, et al. Full genome viral sequences inform patterns of sars-cov-2 spread into and within israel. *Nature communications*, 11(1):1–10, 2020.

F. Obermeyer, M. Jankowiak, N. Barkas, S. F. Schaffner, J. D. Pyle, L. Yurkovetskiy, M. Bosso, D. J. Park, M. Babadi, B. L. MacInnis, J. Luban, P. C. Sabeti, and J. E. Lemieux. Analysis of 6.4 million sars-cov-2 genomes identifies mutations associated with fitness. *Science*, 2022. doi: 10.1126/science.abm1208. URL `https://www.science.org/doi/abs/10.1126/science.abm1208`.

R. B. O'Hara, M. J. Sillanpää, et al. A review of bayesian variable selection methods: what, how and which. *Bayesian analysis*, 4 (1):85–117, 2009.

A. Rambaut, E. C. Holmes, Á. O'Toole, V. Hill, J. T. McCrone, C. Ruis, L. du Plessis, and O. G. Pybus. A dynamic nomenclature proposal for sars-cov-2 lineages to assist genomic epidemiology. *Nature microbiology*, 5(11):1403–1407, 2020.

H. Ritchie, E. Mathieu, L. Rodés-Guirao, C. Appel, C. Giattino, E. Ortiz-Ospina, J. Hasell, B. Macdonald, D. Beltekian, and M. Roser. Coronavirus pandemic (covid-19). *Our World in Data*, 2020. https://ourworldindata.org/coronavirus.

J. M. Smith and J. Haigh. The hitch-hiking effect of a favourable gene. *Genetics Research*, 23(1):23–35, 1974.

Y. Turakhia, B. Thornlow, A. S. Hinrichs, N. De Maio, L. Gozashti, R. Lanfear, D. Haussler, and R. Corbett-Detig. Ultrafast sample placement on existing trees (usher) enables real-time phylogenetics for the sars-cov-2 pandemic. *Nature Genetics*, 53(6): 809–816, 2021.

G. Zanella and G. Roberts. Scalable importance tempering and bayesian variable selection. *Journal of the Royal Statistical Society: Series B (Statistical Methodology)*, 81(3):489–517, 2019.
